# Translational control as a novel regulator of gradient sensing and chemotropism in yeast

**DOI:** 10.1101/2020.12.13.422562

**Authors:** Rita Gelin-Licht, Patrick J. Conlon, Raman Singh, Rohini R. Nair, Gal Haimovich, Camila Baez, Lihi Gal, Maya Schuldiner, Andre Levchenko, Jeffrey E. Gerst

**Affiliations:** Department of Molecular Genetics, Weizmann Institute of Science Rehovot 7610001, Israel; Department of Biomedical Engineering, Yale University, New Haven, CT USA

**Keywords:** chemotropism, yeast mating pathway, gradient sensing, cell fusion, G-protein coupled receptor, MAP kinase, Scp160, Asc1, Gpa2, Gpa1, ribosomal proteins, ribosome specialization, paralog specificity, translation, yeast

## Abstract

The yeast mating pathway regulates haploid cell fusion in response to pheromone signaling via a mitogen-activated protein kinase (MAPK) cascade that controls directional growth (chemotropism). However, the regulators of chemotropic morphogenesis are ill-defined. By using a non-biased genome-wide screen, we identified hundreds of genes that affect mating. An additional screens identified and validated >20 novel positive and negative regulators of pheromone gradient sensing, chemotropism, shmoo development, and mating. Aside from known regulators of exocytosis and endocytosis, genes involved in translational control downstream of the G-protein-regulated pheromone and filamentous growth MAPK pathways were identified. These include the Scp160 RNA-binding protein and the Asc1, Rpl12b, and Rpl19b ribosomal proteins (RPs). Importantly, we demonstrate that pheromone treatment and G*α* (Gpa1) activation stimulate Scp160 binding to (and inhibition of) Asc1, which acts downstream of glucose-activated G*α* (Gpa2) on the filamentous growth pathway. Moreover, we identify both Rpl12b and Rpl19b as RP paralog-specific positive regulators of translation of mating components, including Scp160. Thus, opposing MAPK pathways may converge at the level of translational control to regulate signaling output.

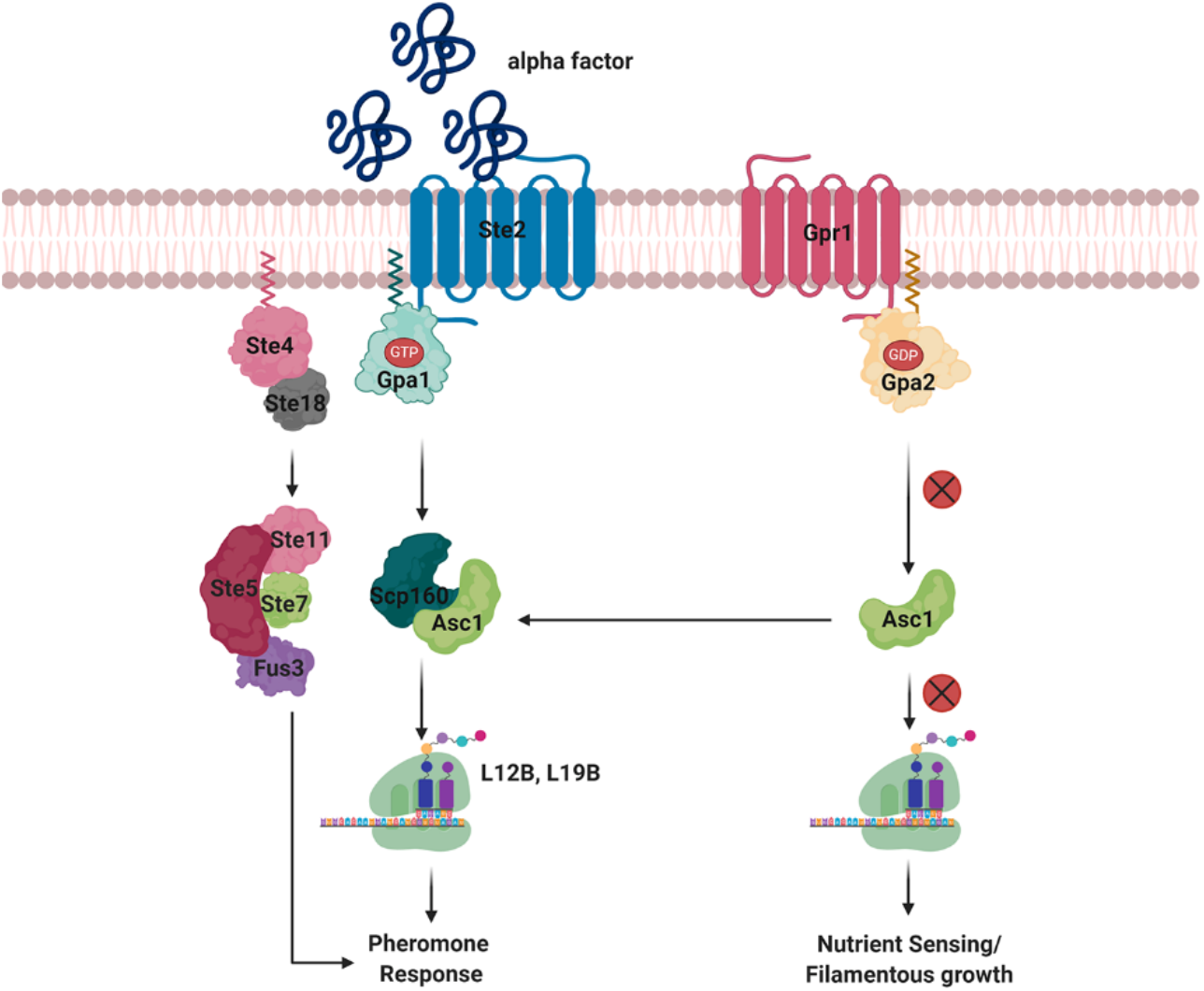

## Introduction

For cells to communicate and coordinate activities, both single- and multi-cellular systems have evolved signaling networks to transduce and amplify input signals, and transform them into phenotypic responses. Such responses include the ability of cells to grow or migrate towards the source of a signal and support specific cell-cell interactions (*e.g.* cell fusion, axon guidance and connectivity) or confer immune responsiveness. Yet, despite extensive study, these processes are not fully understood and the consequences of inappropriate activation or inhibition of such signaling pathways, as well as the relationship between input signals (*e.g.* hormone^1^, drug^2^, or growth factor^3^) and output signals (*e.g.* synaptic function^4^ or metastatic invasion^5^), including those related to human disease (*e.g.* cancer, neurodegeneration) etiology, have not been fully elucidated.

Both chemotropism (directional growth towards an extracellular stimulus)^6–9^ and chemotaxis (directional migration towards an extracellular stimulus)^10–12^ are responses frequently controlled by G-protein coupled-receptor (GPCR) signaling, leading to the activation of mitogen-associated protein kinase (MAPK) networks^13–15^. The cellular mechanisms that underlie these processes are also not fully characterized. The yeast, *S. cerevisiae*, which senses and responds chemotropically to mating pheromone via MAPK signaling^9, 16^ is an ideal system in which to study chemotropism. Discovery of the mating and other MAPK signaling pathways in yeast led to the identification of homologous signaling networks in mammalian cells, principally due to the high degree of genetic and biochemical conservation.

A pheromone-activated MAPK cascade allows *MAT*a and *MATα* haploid yeast cells to respond to pheromone secreted by the opposite mating type to confer the cessation of cell proliferation, polarized chemotropic growth in the direction of the pheromone gradient, and ultimately cell-cell fusion^17^. Pheromone binding to mating type-specific GPCRs activates the G*α*- subunit (Gpa1) and G*βγ* subunit dimer (Ste4/Ste18) in both mating types^17, 18^. Upon activation, G*α* undergoes GDP-GTP exchange and dissociates from G*βγ* to initiate events that precede mating, including gene transcription, cell cycle arrest, as well as the morphological and cytoskeletal changes necessary for mating^9, 17, 18^. In addition to signaling via G*βγ*, a separate signaling function for G*α*/Gpa1 occurs via Scp160^19^, a KH domain-containing RNA-binding protein (RBP)^20^. We have previously shown that RNA-binding by Scp160 is stimulated by pheromone treatment (or direct Gpa1 activation) and delivers mRNAs encoding components of the pheromone-activated MAPK cascade to the site of mating projection (shmoo) formation^21^. Scp160-dependent mRNA delivery to the shmoo tip occurs via directed cortical endoplasmic reticulum (cER) movement and is necessary for continuous correction of the growth of the mating projection^21^, as temporal and spatial accuracy is required for the successful fusion between partners^9, 16^. Thus, Scp160 contributes to chemotropic growth by targeting specific mRNAs to the site of polarization to mediate both gradient sensing, as well as temporal and spatial growth responses to the gradient^21^.

Given the paucity of information regarding how chemotropic growth is specifically controlled, we screened yeast gene deletion and Decreased Abundance by mRNA Perturbation (DAmP) libraries^22^ to identify genes that when mutated, affect mating and specifically chemotropism. Using this approach, we identified over 20 novel positive and negative regulators of chemotropism. While many are associated with exocytosis and endocytosis, which are important for the direct trafficking of pheromone receptors and agglutinins required for mating, we also identify genes involved in translational control. Importantly, we demonstrate that Scp160 function and its role in chemotropic growth is inhibited by Asc1, a 40S ribosomal protein (RP) that is an ortholog of mammalian RACK1^13, 23^. In yeast, Asc1 is involved in the invasive growth response to nutrient limitation via the G*α* (Gpa2)-activated MAPK cascade^24, 25^. Although earlier work suggested that Asc1 interacts preferentially with the inactive GDP-bound form of Gpa2, it is nevertheless required for invasive growth^26^. Earlier studies also found that Asc1 facilitates Scp160 binding to ribosomes, indicating a role in the translation of Scp160-bound messages^27–29^. In addition, Asc1 associates with proteins directly involved in translation initiation (*e.g.* Nip1, Rgp1)^30, 31^ and is essential for translation initiation at elevated temperatures^32^. Finally, an early genetic screen identified Asc1 protein as a possible negative regulator of the pheromone response^33^, but how it affects mating and chemotropism was not further investigated.

Here we not only identify a bilateral relationship between Scp160 and Asc1, we show that Scp160 physically binds Asc1 upon pheromone signaling, but not during nutrient signaling. We also demonstrate that specific ribosomal protein (RP) paralogs are required for the translation of proteins involved in chemotropism and mating. Thus, this study shows that chemotropism is directly regulated by translational control and that the different MAPK pathways (*e.g.* pheromone-activated mating pathway and nutrient-activated filamentous growth pathway) buffer each other’s responses to upstream signaling.

## Results

### An unbiased screen for positive and negative regulators of yeast cell mating

Deficiencies in yeast cell mating can result from a number of reasons, including: an inability to initiate polarized growth in response to pheromone; failure to sense the pheromone gradient; and defects in either cell wall remodeling or cell-cell fusion^34–36^. To select for mutants specifically defective in gradient sensing and chemotropism, we developed a 2-step genetic screening approach to first identify strains deficient in mating and then, specifically, those defective in chemotropic growth.

We first employed high-throughput screening of yeast mutant gene libraries to identify genes involved in mating *per se.* To do this we devised an imaging-based approach to systematically evaluate the mating efficiency for individual gene mutants. The working principle involved mating yeast mutant strains to wild-type (WT) cells and employing image-based detection of both haploids (*i.e.* non-mated cells) and fused zygotes (*i.e.* mated cells) to quantify the overall mating efficiency of the mutant (Figure 1A). To apply this in a high-throughput format, we utilized a pinning robot to mate a WT strain in high density format^37^ to gene knockout^38^ and hypomorphic allele (DAmP)^22^ libraries, which automated haploid cell mixing, mating, and imaging. To increase the visual ability to detect mating events, the knockout library was manipulated to express a cytosolic mCherry reporter (Figure 1A). In order to assess mating efficiency, the *MAT*a mCherry library strains were mixed with GFP-expressing *MATα* WT cells on agar, after which the mating mixtures were collected, fixed, and imaged. After imaging at the mCherry and GFP channels, the number of zygotes (*i.e.* GFP+, mCherry+) and unmated (*i.e.* GFP-, mCherry+) cells were counted, and a mating fraction metric was used to assess mating efficiency for each mutant. Overall, the mating efficiencies of 5,129 mutants (4349 knockouts, including 56 duplicates and 780 DAmP mutants of essential genes) were evaluated in the screen out of the 6221 total genes in these libraries (Figure 1B, Table S1 – Primary screen). We note that 954 mutants (excluding duplicates) were missing from the screen (Tables S1 – Missing). These included yeast strains deleted for genes vital for mating (*i.e. STE* genes, as well as *SCP160*) or cell growth, as the inability to mate or proliferate prevented these strains from completing the first step (*i.e.* forming zygotes containing the mCherry gene) of the library creation (Fig 1A, top). This list also includes 146 genes that were missing for technical reasons. These strains were added in later steps of the screen.

**Figure 1.**
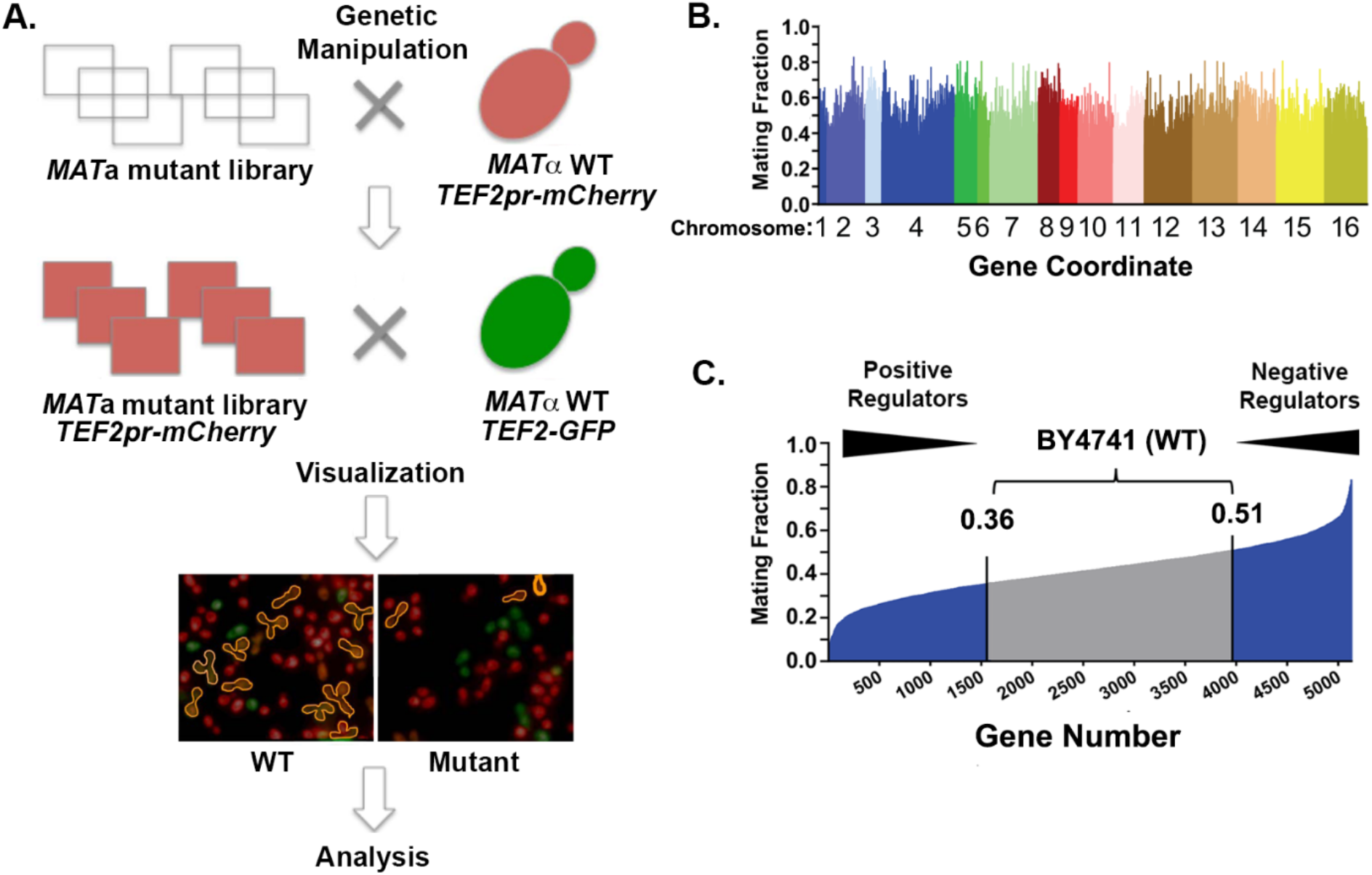
Systematic approach for and identification of regulators of yeast cell mating. (A) Non- biased screen for genes required for yeast cell mating. Yeast mutant libraries (knock-out and DAmP) were crossed against a WT query strain carrying a cytosolic marker fused to mCherry using the synthetic gene array (SGA) methodology. *MAT*a cells derived from the modified knock-out and DAmP libraries were then mixed with a WT *MATα* strain that expresses a cytosolic marker fused to GFP for 3hrs. The crosses were then automatically imaged for GFP, mCherry, and mixed pattern fluorescence detection, followed by analysis. (B) Mating fraction analysis. Calculation of the mating fraction (*i.e.* percentage of cells that underwent mating) for each *MAT*a strain was determined. After thresholding and scoring, the mating fraction for each strain was calculated as the number of diploid cells (GFP/RFP) divided by the number of diploid (GFP/RFP) and haploid (RFP + GFP) cells together. After normalization (using the average mating efficiency for each plate), the mating fraction score for each mutant was plotted. A score of 1.0 designates that 100% of the *MAT*a cells underwent mating. The plot shows the data for 5129 yeast mutants tested and illustrated in color-code across the different yeast chromosomes, as listed. (C) Sorted mating fraction scores. Shown are the normalized mating fraction scores for the *MAT*a mutants sorted from lowest to highest fraction. The gene number is listed as in column A of Table S1 – *primary screen*. Mating fractions were normalized by the average mating efficiency for each plate. The grey region indicates the range of mating fraction scores obtained using WT BY4741 cells. Mutants whose scores fell within this range (0.36- 0.51) were not considered for further analysis.

Image analysis of the mutants (Table S1; sheet – *Primary screen*) suggested that ∼2770 genes may either regulate or affect mating efficiency, based upon the fraction of cells that underwent mating, compared to wild type (WT) BY4741 cells that show a range in mating from 0.36-0.51 of the population (Figure 1C). Out of this long list of genes that affect mating, we selected a list of 581 genes (Table S1, sheet – *Second screen*). We excluded many genes that we suspected might not be specific to mating (*e.g.* genes encoding mitochondrial and other organelle-specific functions, as well as genes involved in general transcription, translation, and metabolic functions). Initially, we also excluded general translation factors (e.g. ribosomal proteins). However, we return to some of these genes later (see below). Additionally, we included the 146 genes that were missing from the primary screen due to technical reasons.

To validate the reliability of the primary screen, a second automated screen was performed with this select group of 581 genes applying the same conditions (Figure 2A, Table S1, sheet – *Second screen*). We found that screen reproducibility was high: 65% (379 genes) of positive or negative regulators from the first screen had similar qualitative results in the second screen relative to the controls. Moreover, 86% (327 genes) of the genes that were qualitatively reproducible also exhibited quantitative reproducibility, within a range of ±0.1 of the primary screen’s mating fraction (Fig. 2A, Table S1, sheet – *Second screen*). Of the additional 146 genes examined, 6.2% (9 genes) appeared to be positive regulators of mating and 74.5% (108 genes) negative regulators, whereas 19.3% (28 genes) scored in the WT range (Figure 2A).

**Figure 2.**
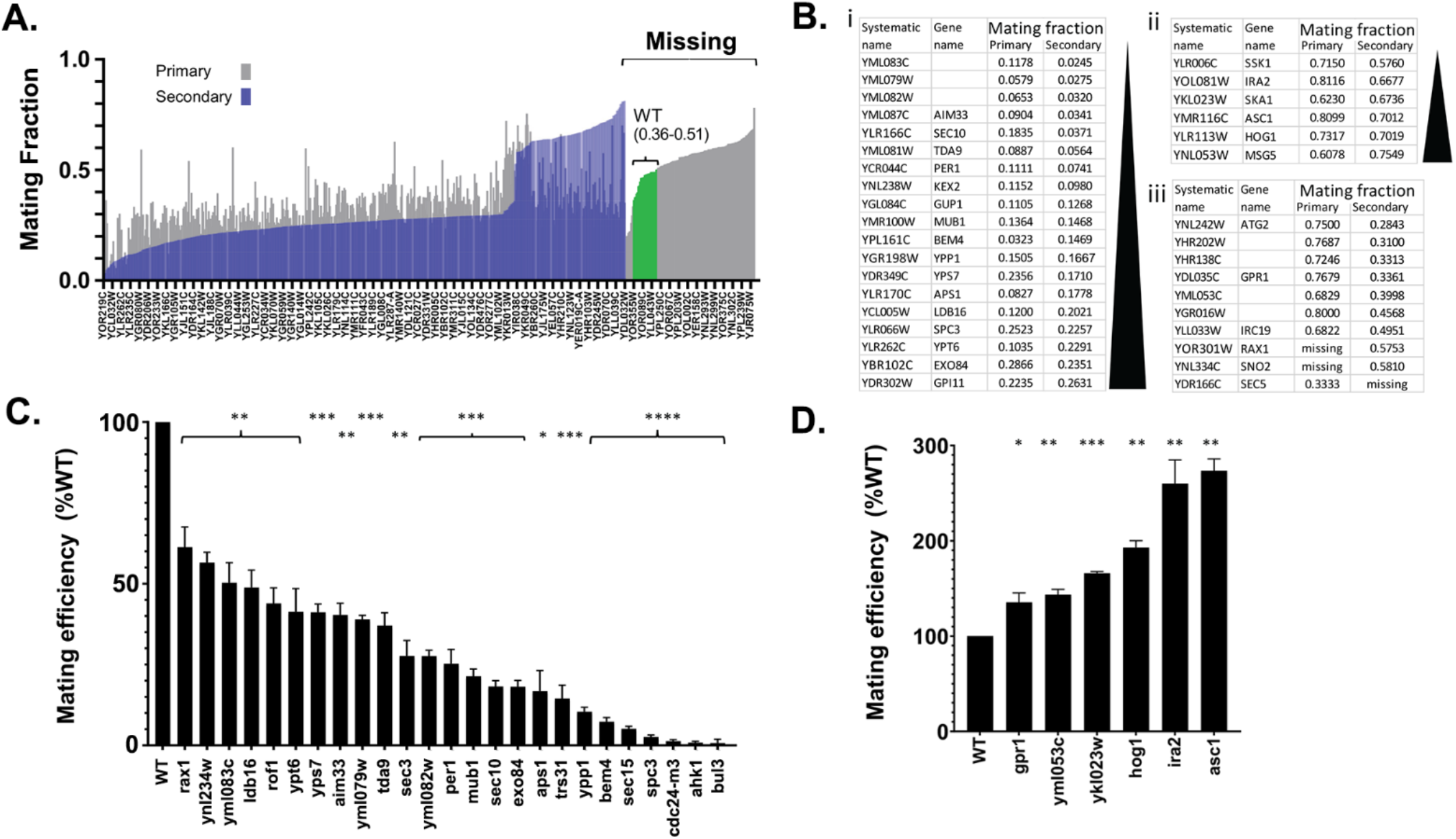
Candidate selection and validation of mutants from the mating screen. (A) Sorted mating fraction scores. Shown are the normalized mating fraction scores for the *MAT*a mutants sorted from lowest to highest fraction. The given systematic gene names are listed as in column F of Table S1 – primary screen. Purple – results of primary screen. Gray – results of secondary screen. Green - Mutants whose scores fell within the WT range. *Missing* – strains absent from primary screen. (B) Partial lists of mating-defective (upon mutation) genes identified from both screens is given, with those having the lowest mating efficiency shown on the top. (i) Genes encoding positive regulators of mating. (ii) Genes encoding negative regulators of mating. (iii) This list includes strains with only one score or with inconsistent scores. (C, D) Manual validation of positive regulators (C) and negative regulators (D). Quantitative mating assays were performed on the most positive regulators using manually constructed deletions/mutations to validate results of the screen. The Y axis shows the % of mating, with the mating efficiency of control WT cells (BY4741) designated as 100%. The *cdc24-m3* allele was included as a control. Gene names listed on the X axis indicate the mutant genes tested. Shown are the average value of at least three replicates ± standard deviation. * indicates *p* ≤0.05; ** indicates *p* ≤0.01; *** indicates *p* ≤0.001; **** indicates *p* < 0.0001

Mutants of known positive regulators (*i.e.* genes known to be important for promoting mating) showed severe mating defects (*e.g.* scoring at the bottom 10% of mating efficiency). These included known genes such as *AGA2, AXL1, STE2, STE24, STE50*, *FUS1*, *FUS3*, *BAR1, BEM4* and *CDC42* (Table S1, sheet – *Second screen*). The list of positive regulators also included previously characterized genes, like those encoding components of the exocyst (*e.g. SEC10, EXO70* and *EXO84*) or other components of the secretion pathway (*e.g*. *SEC1* and *SEC26*) that are involved in secretion, but were not known to mediate mating *per se*^9, 39, 40^. In contrast, mutants of known negative regulators (*e.g.* genes known to be antagonistic to the mating pathway) showed highly proficient mating (*i.e.* top 5% of mating efficiency). These included genes such as *HOG1*, *PBS2*, and *MSG5,* (Table S1, sheet – *Second screen*). *HOG1* and *PBS2* encode proteins in MAPK cascade involved in the cellular response to high osmolarity conditions^41^. Msg5 is a phosphatase involved in Fus3 regulation and inhibits its nuclear accumulation^42, 43^. In addition to these known (or more obvious) genes, others were identified that have not been described in the context of mating, including uncharacterized ORFs identified as negative regulators (*e.g. YKL222C* and *YKR045C*), and positive regulators (*e.g. YGR237C, BRP1, FYV6* and *YPR114W*), as well as nine genes that occupy a common region in Chr. XIII (*YML089C, YML087C, YML086C, YML084W, YML083C, YML082W*, *YML081C*, *YML080W* and *YML079W*). From these validated results, we further evaluated potentially interesting positive and negative regulators of mating. We performed manual deletion of non-essential genes and used a tetracycline-repressible (tet-off) promoter system^44^ for essential genes. The list includes 56 genes, which were selected by various criteria (Table S1, sheet – *Manual validation,* column F). In addition to genes that were consistent between both screens (which included five of the nine genes on Chr. XIII) (Figure 2B i and ii), we chose genes with a high score in the 1^st^ or 2^nd^ screen (*e.g. RAX1, GPR1, IRC19, YGR016W, SNO2*), since we figured these could benefit from additional validation (Figure 2B iii). We also chose from the “Missing” list several genes that are implicated in mating (*e.g. SCP160, BUL3*), genes related to some of our validated genes (*e.g. SCP160, SRO9, EGD2, SYG1, AHK1* and exocyst subunits), as well as ten randomly chosen genes, since we suspected that many of the “Missing” genes are defective in mating.

Manual validation was first used to verify the mating phenotype of the query strains. We performed quantitative mating assays in crosses against WT cells (Figure 2C and D, Table S1 - *Manual validation,* column G). We also included a negative control, *cdc24-m3*, which contains a recessive mutation in the exchange factor for the Cdc42 GTPase and is severely inhibited in chemotropism and mating^45^. The results obtained from manual manipulation were in high qualitative agreement with the results of the library screen (*i.e.* 29 of 35 genes, Figure 2C and Table S1) and showed a varying degree in the regulation of mating. This was true for both positive (Figure 2C) and negative regulators (Figure 2D) of mating. The remaining 6 mutants included 3 that displayed a mating efficiency similar to WT and 3 that showed the opposite mating efficiency than expected from the previous screen. Among mutants not present in the initial screens, but examined manually, we found that 18 genes positively regulated mating, while 2 genes (both from the randomly chosen list) behaved as wild-type cells and one gene exhibited negative regulation of mating (Figure 2B and D and Table S1, sheet – *Manual validation, Columns G and H*).

### A screen for positive and negative regulators of chemotropism

To assess the specific contribution of candidate genes to chemotropism, we performed an additional type of screen called the “pheromone confusion” assay^46^. The confusion assay compares the mating efficiency of *MAT*a and *MATα* haploid cells either with or without exogenous *α*-factor added to the mating mixture (Figure 3A). The mating efficiency of WT cells is significantly reduced when exogenous *α*-factor is present because WT *MAT*a cells are no longer able to detect the pheromone gradient coming from their *MATα* partner. However, mutant *MAT*a cells that have a defect in chemotropism do not show any further reduction in mating efficiency when the gradient is disrupted by added *α*-factor.

**Figure 3.**
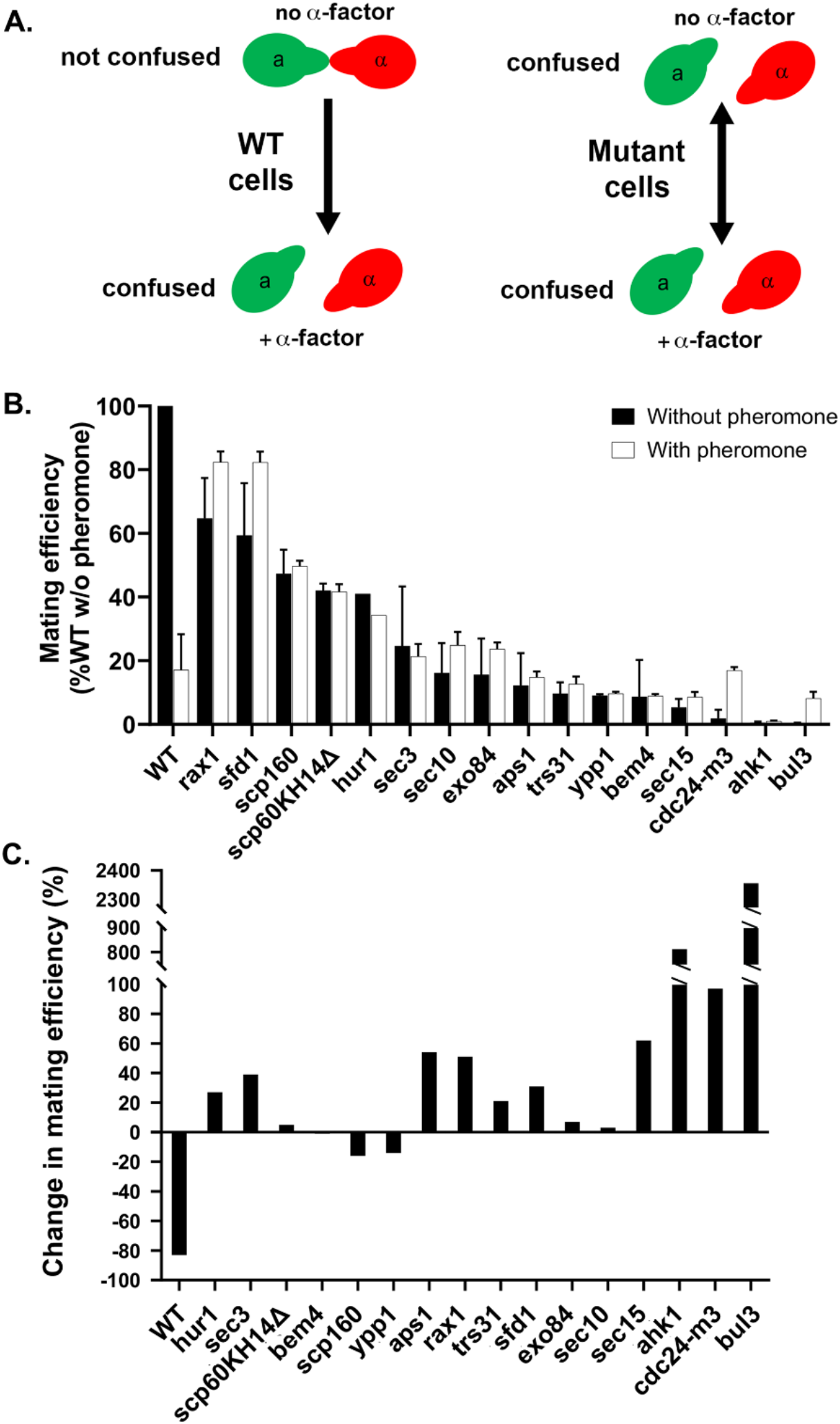
Candidate selection and validation of mutants from the mating screen using the confusion assay. (A) Schematic of the pheromone confusion assay. Left side: WT *MAT*a and *MATα* cells normally undergo mating (*not confused*) in the absence of added *α*-factor. However, when mixed in the presence of exogenously added pheromone at saturating levels they are unable to undergo directional growth towards their mating partner and fail to mate (*confused*). Right side: *MAT*a mutants of the positive regulators identified in the screen are already confused (*confused*) and, thus, show no further inhibition in mating to *MATα* WT cells upon the addition of exogenous pheromone (*confused*). This indicates the relevance of the corresponding gene in chemotropism. (B) Validation of chemotropism defects using the confusion assay. Confusion assays were performed on mutants previously indicated as positive regulators. Positive controls included *SCP160* lacking its KH domain (*scp160KH14Δ*), as well as *cdc24-m3*. Common names listed on the X axis indicate mutant genes tested. In all cases, *p* ≤0.0001. (C) Measure of the change in mating efficiency in cells treated with excess pheromone. The difference in mating efficiency between cells treated with exogenous pheromone (+*α*-factor) and untreated cells (shown in *B*) was calculated as % change and plotted as a histogram. In all cases, *p* ≤0.0001.

To evaluate their chemotropic properties, we determined the mating efficiency of 40 manually validated positive regulators either with or without added pheromone (Figure 3B and Table S1, sheet – *Manual validation, columns I-J*). We did not examine deletions of the negative regulators of mating since they are already likely to be indifferent to external pheromone. We found 18 mutants to be highly deficient in chemotropism in the confusion assay, since they exhibited <20% decrease in mating efficiency or in most cases a significant increase therein, upon exposure to exogenous pheromone at saturating concentrations, relative to untreated cells (Figure 3B and C). We identified this group of mutants as novel chemotropism-related candidates (*i.e. sec3Δ, sec10Δ, sec15Δ, exo84Δ, trs31Δ, ypp1Δ, bul3Δ, ahk1Δ, bem4Δ, sfd1Δ, rax1Δ, sro9Δ, sno2Δ, spc3Δ, ldb16Δ, aps1Δ, hur1Δ* and *scp160Δ*). This group also includes controls such as *scp160KHΔ14*, an *SCP160* allele which lacks the last KH RNA-binding domain of Scp160 and is thought to be unable to bind RNA^47^, and the *cdc24-m3* allele. In contrast, the mating efficiency of WT control cells, as well as 22 mutants, was greatly inhibited by the exogenous addition of pheromone (Figure 3B and Table S1, sheet – *Manual validation, Columns I-K)*. Interestingly, all five genes from Chr. XIII showed a mating, but not chemotropism, defect. It remains to be determined why genes in this region are defective in mating.

### Quantitation of MAPK activity and shmoo formation

To help understand how these different genes affect chemotropism and mating, we quantified both the signaling and morphological responsiveness to pheromone stimulation of both the positive regulators listed above and negative regulators of mating (*hog1Δ, ygr016wΔ, ska1Δ, irc19Δ, gpr1Δ, msg5Δ, ira2Δ, asc1Δ*). We determined the relationship between defects in gradient sensing and activity of the MAPK pathway in the different strains using Fus1-GFP as a reporter^17, 48, 49^. In parallel, we examined shmoo formation as an indicator of readout at the morphological level. By employing flow cytometry, Fus1-GFP expression levels (Figure 4A) and shmoo projection formation (Figure 4B) were assayed. Mutants in positive regulators (*e.g. scp160Δ*, *sec15*, *bul3Δ*, *ahk1Δ*) demonstrated reduced Fus1 expression (*e.g.* 0.9% up to 60% of WT) in parallel with poor shmoo formation (*e.g.* 3% up to 66%) in all cases (Figures 4A and B). Positive controls in this experiment included the *cdc24- m3* and *scp160KHΔ14* alleles. However, mutants in the negative regulators of mating exhibited mild to no effect on MAPK pathway reporter expression compared to WT cells (*e.g.* 73% to 120% of WT), but showed somewhat lower levels of shmoo formation (*e.g.* 40% up to 75% of WT). Thus, mutants in the negative regulators, which show enhanced mating (Figure 2D), do not show enhanced signaling or shmooing.

**Figure 4.**
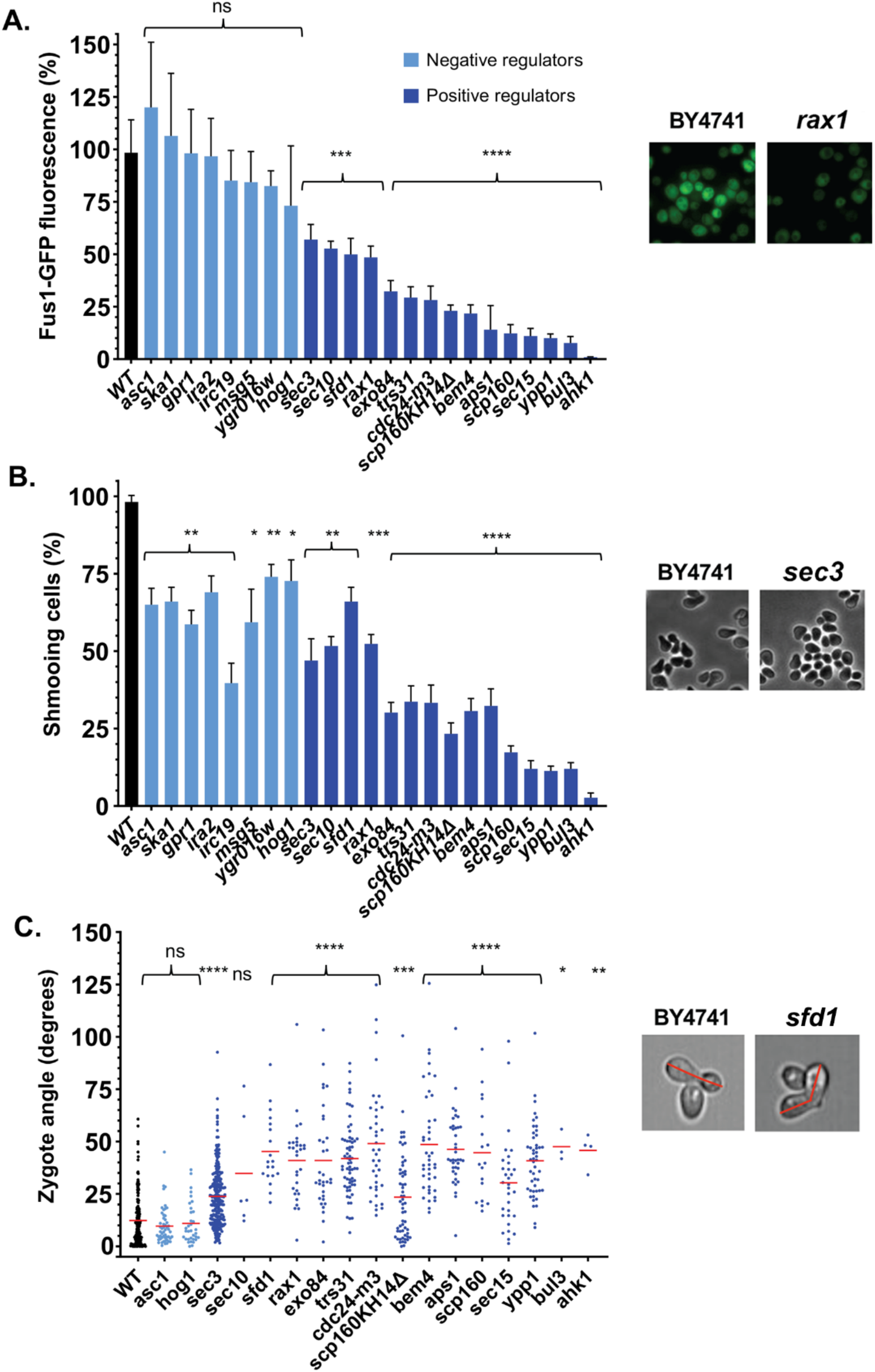
Relationship between activity of the MAPK pathway, cell morphology, and defects in gradient sensing. (A) Measurement of MAPK activity in the positive regulators of chemotropism and negative regulators of mating. Reporter activity in mutants of negative regulators of mating and positive regulators that show the “confusion” phenotype (Figure 3). The photos illustrate the activity of WT (BY4741) cells versus one of the mutants, as an example. (B) Measurement of shmoo formation in the positive regulators of chemotropism and negative regulators of mating. WT and mutant *MAT*a cells were grown and treated for 2hrs in the presence of 5μM α-factor at 30°C, prior to scoring for mating projection (shmoo) formation. The photos illustrate the shmooing of WT (BY4741) cells versus one of the mutants, as an example. (C) Measurement of the angle of zygote formation in the positive regulators of chemotropism and negative regulators of mating. Each dot in the graph represents a single zygote. The photos illustrate the zygote angle of WT (BY4741) cells versus one of the mutants, as an example. For all panels, gene names listed on the X axis indicate the mutant genes tested. ns – not significant; * indicates *p* ≤0.05; ** indicates p≤0.01; *** indicates p≤0.001; **** indicates *p* ≤0.0001.

### Quantitation of the angle upon zygote formation

To assess whether the individual mutants show *bona fide* changes in chemotropism, we applied high-throughput imaging to mixed mutant *MAT*a and WT *MATα* cell populations. The method is based on the observation that the angle between two mating cells during zygote formation depends upon their precise orientation towards one another^50^. If their orientation (*i.e.* based upon gradient sensing) is optimal and cells grow directly towards one another, their angle of fusion is small and close to 0°. In contrast, an imperfect orientation to the pheromone gradient exhibits large fusion angles^50^.

By using an ImageStream flow cytometer, we found that mutants of genes identified in the screen as positive regulators of mating demonstrate a profound defect in chemotropism (*i.e.* the average angle of orientation is ∼40.3°<u>±</u>2.2° (average<u>±</u>SEM) for mutant crosses against WT cells versus 12.4°<u>±</u>1.0° (average<u>±</u>SEM) for WT crosses against WT cells (Figure 4C). In contrast, the negative regulators of mating exhibited an improved orientation, being even slightly better than WT cells (*i.e.* the average angle of orientation is 10.2°<u>±</u>1.0° (average<u>±</u>SEM) for mutants).

### Categorization of chemotropism mutants

Based upon the results from all of the various assays performed, we categorized the mutant strains according to their quantified similarity in phenotype, with respect to mating MAPK signaling, shmoo formation, and zygote angle. As mentioned above, many of the genes identified in the screen encode proteins with known functions connected to exocytosis and endocytosis, which regulate both pheromone receptor surface expression as well as the localization of polarity patches involved in gradient sensing and shmoo formation^9^. For convenience, we categorized the various gene hits into four groups.

**Group I** genes were characterized as positive regulators of mating having ∼50% of MAPK signaling activity and ∼50-65% shmoo formation capacity, as compared to WT. The zygote angle degree ranges between 24-45°. These genes include two subunits of the exocyst complex (*SEC3* and *SEC10*), which plays a critical role in vesicle tethering and secretion^51^. This group also includes *RAX1*, which encodes a protein required for bud/shmoo site selection and cell polarization, and whose deletion affects placement and angle of the mating projection and is thought to control Gpa1 and Cdc24 function^52^. The last member of this group is an uncharacterized gene (*SFD1*).

**Group II** genes are also positive regulators and show a more severe phenotype, with 15-30% MAPK signaling activity, 30-35% shmoo formation, and >40° zygote angle. This group includes the exocyst complex factor *EXO84*, a gene encoding the small subunit of the clathrin-associated adaptor complex, *APS1*^53^, which likely functions in Ste2 internalization, and *TRS31*, a core component of the TRAPP vesicle tethering complex that regulates traffic to and from the Golgi, but also facilitates post-Golgi vesicle trafficking^54^. This group also includes Bem4, which is a regulator of Cdc24 (and, thus, Cdc42 activation) and also interacts with the Ste11 MAPK in control of the mating signal^55^. The positive control strain, *cdc24-m3*, also belongs to this group.

**Group III** mutants are positive regulators that have the most severe phenotype, showing 1-12% of Fus1-GFP expression and 3-17% shmoo formation compared to WT. However, the zygote angle ranges similar to Group I (*e.g.* 30-47°). This group also includes trafficking-related factors such as *SEC15*, which encodes another exocyst complex subunit, and Ypp1, which is required for Stt4 phosphotidylinositol-4 kinase localization to the plasma membrane^56^ and the clathrin-dependent endocytic internalization of Ste2^57^. *AHK1* encodes a putative scaffold component of the Hkr1 sub-branch of the HOG signaling pathway and is required for transmission of the osmolarity stress response^58^, as well as other signals via the Ste11 MAPK, which is required for mating. *BUL3* encodes a protein involved in the ubiquitin-mediated sorting of plasma membrane proteins associated with Rsp5 ubiquitin ligase activity^59^, which mediates pheromone receptor internalization^60^. The last member of this group is *SCP160*, an RNA binding protein previously shown to have a direct role in mating and chemotropism^21^. Interestingly, *scp160KH14Δ*, which lacks RNA-binding activity, shows a phenotype in between Groups II and III. Thus, it is possible that this mutant retains some mating functions unrelated to RNA-binding. Notably, there is no clear correlation between mating efficiency (Figure 2, Table S1) or the difference after confusion of the different strains (Figure 3B), to the grouping based on the above three assays. Thus, multiple factors may possibly affect mating efficiency and the confusion effect for each mutant.

Arguably, many of Group I-III genes are likely involved in pheromone receptor transport and recycling and, thus, might be predictable in their defects in chemotropism and mating. However, **Group IV** genes are negative regulators of mating (*e.g. YGR016W, SKA1, IRC19, GPR1, IRA2, MSG5, HOG1* and *ASC1*), whose mutants demonstrate similar to or slightly less than WT level of MAPK signaling (73-120%), and the two mutants we tested show an optimal zygote orientation, slightly better than WT. However, these mutants still have defects in shmoo formation (*e.g.* 40-75% of WT), yet show enhanced mating efficiency in ways that may be distinct from receptor recycling mechanisms. These genes comprise the negative regulators of mating and many appear to be involved in MAPK cascades. *GPR1* encodes the GPCR involved in glucose sensing in the filamentous growth pathway and acts upstream of adenylyl cyclase in a Ras-independent fashion^61, 62^. *IRA2* encodes a Ras-GTPase activating protein that lowers the activity of the Ras pathway and adenylyl cyclase^63^. *HOG1* encodes a MAPK involved in osmosensing^64^. *MSG5* encodes a phosphatase involved in Fus3 regulation and inhibits its nuclear accumulation^42, 43^. Finally, *ASC1* encodes a ribosomal protein that acts downstream of the Gpr1 signaling to confer filamentous growth^13, 23, 26^ and facilitates Scp160 association with polysomes^29^. Thus, five of the negative regulators of mating are directly involved in signaling along the filamentous growth pathway or the pheromone signaling pathway.

Of the others, *SKA1* encodes a protein proposed to facilitate exosome (3’-5’ exonuclease) function in the catabolism of non-ribosome-associated mRNAs^65^, while *YGR016W* encodes a protein of unknown function. While the connection of these proteins to the negative regulation of mating is unknown, both proteins were shown to be substrates of the E3 ubiquitin ligase, Rsp5^66^. Finally, *IRC19* has been suggested to function with other genes in restricting recombination events to regulate cell division^67^.

### Deletion of *ASC1* enhances Scp160-mediated RNA binding and mating

The positive role of Gpa1-activated Scp160 in chemotropism and mating has been largely defined^19, 21^, and is validated here (Figure 3B). However, we also identified Asc1 as a negative regulator of mating (Figure 2C and D). Asc1 is a ribosome-associated protein that facilitates Scp160 binding to polysomes^29^ and acts as a downstream effector of Gpr1- and Gpa2-mediated filamentous growth^26^. This suggests that Scp160 and Asc1 functions are antagonistic to each other, and may illustrate how the two MAPK pathways intersect to regulate signal transduction at the level of translational control.

Since the RNA-binding function of Scp160 is specifically activated by pheromone signaling or constitutively active Gpa1 (Gpa1^Q323L^)^21^, we examined whether the deletion of *ASC1* has an effect upon the binding of mRNAs (Figure 5). We expressed FLAG epitope-tagged Scp160 from the genome along with either native or activated Gpa1 from single-copy plasmids in WT or *asc1Δ* cells. In these cells, we examined the binding of Scp160 to an mRNA (*FUS3*) known to bind upon activation and another (*ASH1*) that does not^21^. As expected, the expression of Gpa1^Q323L^ enhanced *FUS3* (42-fold) but not *ASH1*, mRNA binding to FLAG-Scp160 in WT cells, whereas the expression of non-activated Gpa1 had no effect (Figure 5). Interestingly, the deletion of *ASC1* led to increased *FUS3* binding to FLAG-Scp160 in WT cells (either with or without the expression of native Gpa1), and strong enhancement (∼200-fold binding) in cells expressing activated Gpa1 (Figure 5). Thus, Asc1 expression inhibits the RNA-binding function of Scp160, particularly in the absence of pheromone signaling.

**Figure 5.**
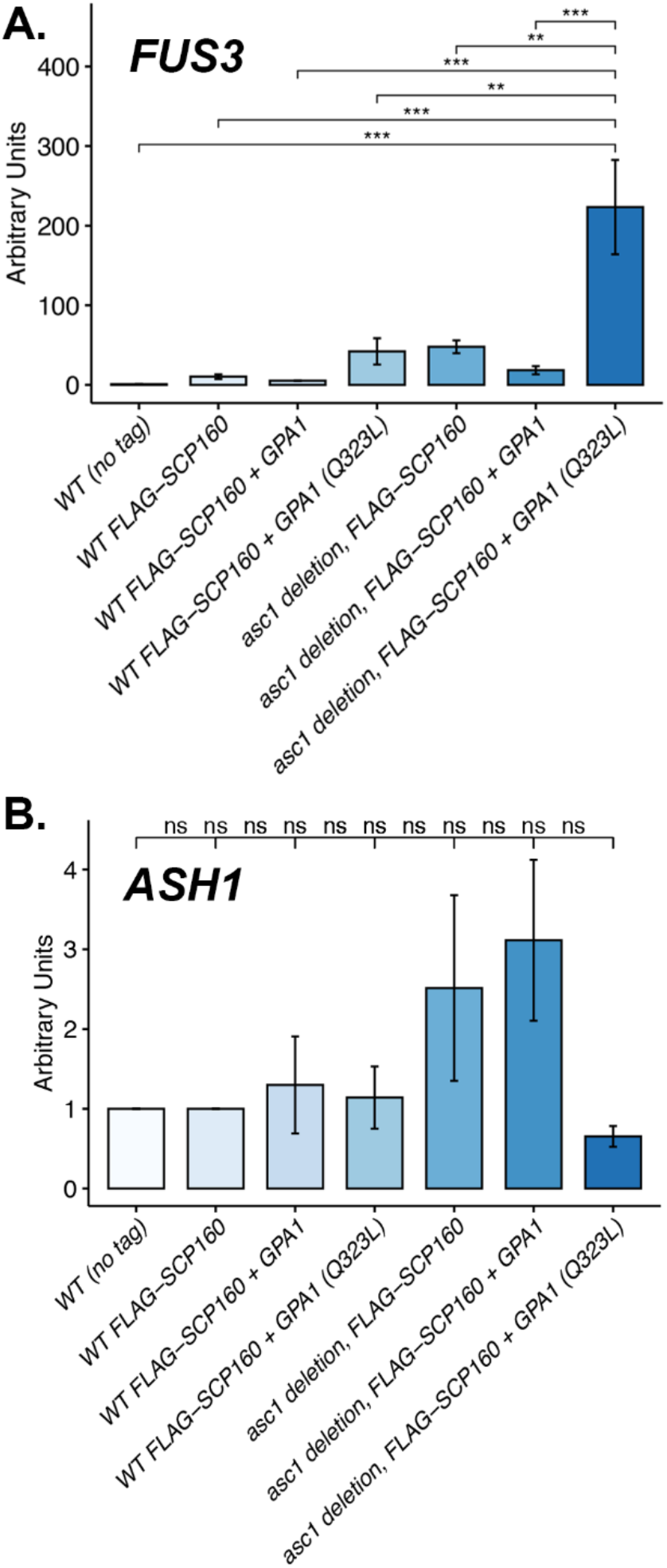
*FUS3* mRNA binding by Scp160 is modulated in an *ASC1*-dependent manner. Either wild-type (WT) cells or *asc1Δ* cells expressing FLAG epitope-tagged Scp160, both FLAG-Scp160 and Gpa1^Q323L^, or FLAG-Scp160 and native Gpa1, as well as WT control cells, were grown to mid-log phase, lysed, and incubated with anti-Flag M2 affinity gel to immunoprecipitate Scp160. RNA was extracted from the TCL samples and immunoprecipitates, and used as a template for real-time qPCR. Specific primer pairs were used to detect the *FUS3* (upper graph) and *ASH1* (lower graph) mRNAs. ns – not significant; ** indicates *p* ≤0.01; *** indicates *p* ≤0.001. Note that the levels of Scp160- FLAG protein in TCL samples are not shown since these were below the level of detection.

Since the deletion of *ASC1* stimulates mating (Figure 2D), we determined whether the over- expression of *ASC1* in cells inhibits mating. We examined the mating efficiency of WT cells overexpressing *ASC1* from a multicopy plasmid, in comparison to WT cells alone or over-expressing *SCP160*, or to the *scp160Δ* and *asc1Δ* deletions (Figure S1). We found that the deletion of *ASC1* led to a 4.2-fold increase in mating efficiency, while its over-expression resulted a 24.4-fold decrease (Figure S1A). In contrast, the deletion of *SCP160* led to a 4.4-fold decrease in mating and its overexpression had almost no effect. Thus, both *ASC1* over-expression and deletion had robust effects on mating, whereas that of *SCP160* was more limited. Since *ASC1* has an intron, we also examined whether intron presence or removal had an effect on its ability to inhibit mating. However, both intronless and native *ASC1* had similar inhibitory effects (Figure S1B).

### Scp160 binds Asc1 upon pheromone signaling

Given that Asc1 interacts with Scp160^29^ and acts as both an inhibitor of the mating pathway (Figure 2D) and modulator of the RNA-binding function of Scp160 (Figure 5), we examined whether its interaction with Scp160 is regulated via the Gpa1-mediated pheromone signaling pathway or the Gpa2-mediated filamentous growth signaling pathway (Figure 6). We expressed HA epitope-tagged Asc1 in WT or FLAG-Scp160 cells and co-expressed either Gpa1^Q323L^ or activated Gpa2 (Gpa2^Q355L^), and performed pulldowns using anti-HA antibodies. We observed that HA-Asc1 precipitated little Scp160 alone, but showed a large increase in Scp160 binding in the presence of added exogenous pheromone (*α*-factor) or Gpa1^Q323L^ (*i.e.* 4.4-fold and 3.5-fold, respectively) (Figure 6). In contrast, no Scp160 binding to Asc1 was observed in cells expressing Gpa2^Q355L^. Thus, Scp160 binds to Asc1 upon pheromone stimulation, but not during filamentous growth signaling.

**Figure 6.**
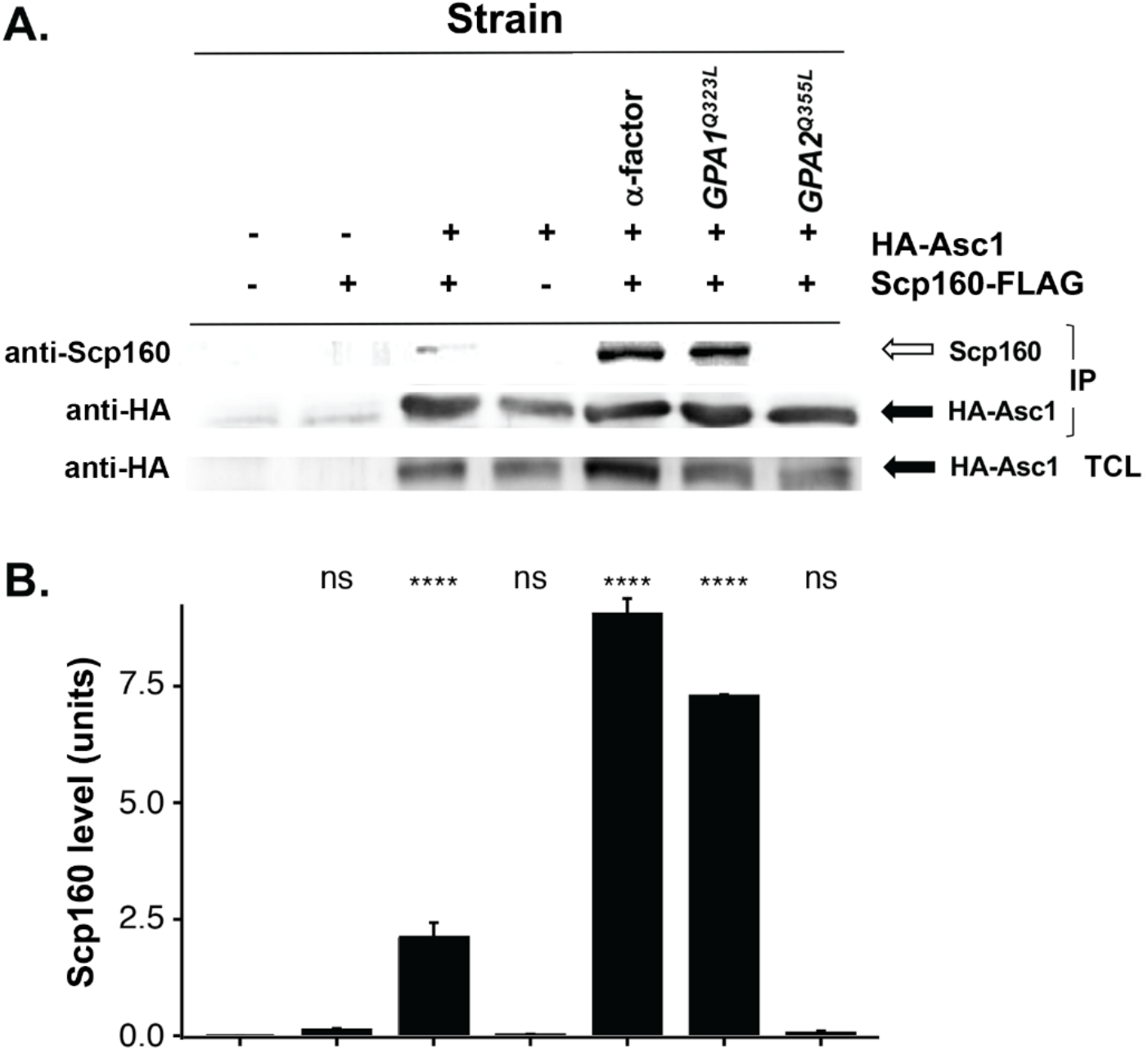
Scp160 binds Asc1 in a pheromone-activated manner. (A) Cells co-expressing HA-tagged Asc1 and Scp160-FLAG were left either untreated or treated with pheromone (5μM α-factor), or expressed Gpa1^Q323L^ or Gpa2^Q355L^ from single-copy plasmids, were used to examine the effect of pheromone pathway activation on the interaction between Asc1 and Scp160. Controls included wild-type (no tag) cells and cells expressing either HA-Asc1 or Scp160-FLAG alone. Cells were grown to mid-log phase, lysed, and incubated with anti-HA affinity gel to immunoprecipitate proteins. Samples of each immunoprecipitate (IP) were separated on a 9% SDS-PAGE gel, blotted, and probed with anti-HA and anti-Scp160 antibodies. The filled arrow indicates HA-Asc1; the open arrow indicates Scp160-FLAG. (B) Quantification of the protein bands from (*A*) using ImageJ. A representative experiment (n=3 experiments) is shown.

### *FUS3* mRNA granule formation and localization in pheromone-treated cells depends on both Scp160 and Asc1

We previously demonstrated that *FUS3* mRNA localizes to the tip of the mating projection (shmoo) upon pheromone stimulation^21^. We examined the effect of either *SCP160* or *ASC1* deletion on *FUS3* mRNA granule formation and localization (Figure S2). Using MS2 aptamer-tagged *FUS3* expressed from the genome under its native promoter, we observed that whereas WT and *scp160Δ* cells had mostly single *FUS3* mRNA granules, *asc1Δ* cells had multiple granules in all cells scored (Figure S2A and B). This suggests that Asc1 has a role in the integrity of the *FUS3* granules. Moreover, we noted that the multiple *FUS3* mRNA granules observed in *asc1Δ* cells were not localized to the shmoo tip or body (*i.e.* near the tip), but instead were mislocalized throughout the cell body (Figure S2C). Likewise, the single mRNA granules observed in *scp160Δ* cells were also mislocalized (Figure S2C), thus, both Asc1 and Scp160 are required for the normal trafficking and localization of *FUS3* mRNA.

### Fus3 expression and phosphorylation are not altered in cells lacking either *SCP160* or *ASC1*

Given that the deletion of either *ASC1* or *SCP160* affects the localization of Scp160 target mRNAs, like *FUS3*, we examined their effect on Fus3 protein expression and phosphorylation in response to pheromone signaling. We used anti-phospho ERK antibodies that recognize both Kss1-P (at 43kDa) and Fus3-P (at 41kDa) upon pheromone treatment^68^ to detect the proteins in cell lysates (Figure S2D). We observed no differences in the levels of these phosphorylated MAPKs upon the deletion of either gene (Figure S2D). This indicates that pheromone signaling up to this point remains intact.

### Specific paralogs of ribosomal proteins control chemotropism

Given that Asc1 and Scp160 help control protein translation in cells, we re-examined the results of our initial screen (Figure 1B and C, Tables S1 and S2) for genes involved in translation (*e.g.* RPs) that had been removed from further analysis. Specifically, we were curious to see whether there are RP paralogs with opposing phenotypes with regards to mating, since recent discoveries from our lab and others suggest that RP paralogs function differently under distinct conditions^69, 70^. We found that out of 42 paralog pairs present in our primary screen, 20 pairs (48%) had A and B paralogs with dissimilar mating fraction scores (*i.e*. >0.1) and phenotypes (Table S2). Of these pairs, 9 had a mating fraction difference of >0.25. In addition, we found 6 RP genes that were represented by only one paralog in our libraries and showed either a negative or positive regulator phenotype (Table S2). We evaluated the role of 6 selected RPs in mating assays after manual deletion (*e.g. RPL9A*, *RPL12A*, *RPL12B*, *RPL19B, RPS21B,* and *RPS27B*) (Table S2, Figure 7A). In agreement with the primary screen results, the deletions of *RPL12B* or *RPL19B*, but not *RPL12A* (Table S2, Figure 7A), were found to strongly inhibit mating, while those of *RPL9A*, *RPS21B,* and *RPS27B* increased mating (Table S2, Figure 7A). Given that *RPL12B* and *RPL19B* encode positive regulators of mating, we subjected their deletion mutants to the pheromone confusion assay and found that neither showed additional effects upon mating efficiency in the presence of exogenous *α*-factor (Figure 7B). Thus, *rpl12bΔ* and *rpl19bΔ* are suspected chemotropism mutants.

**Figure 7.**
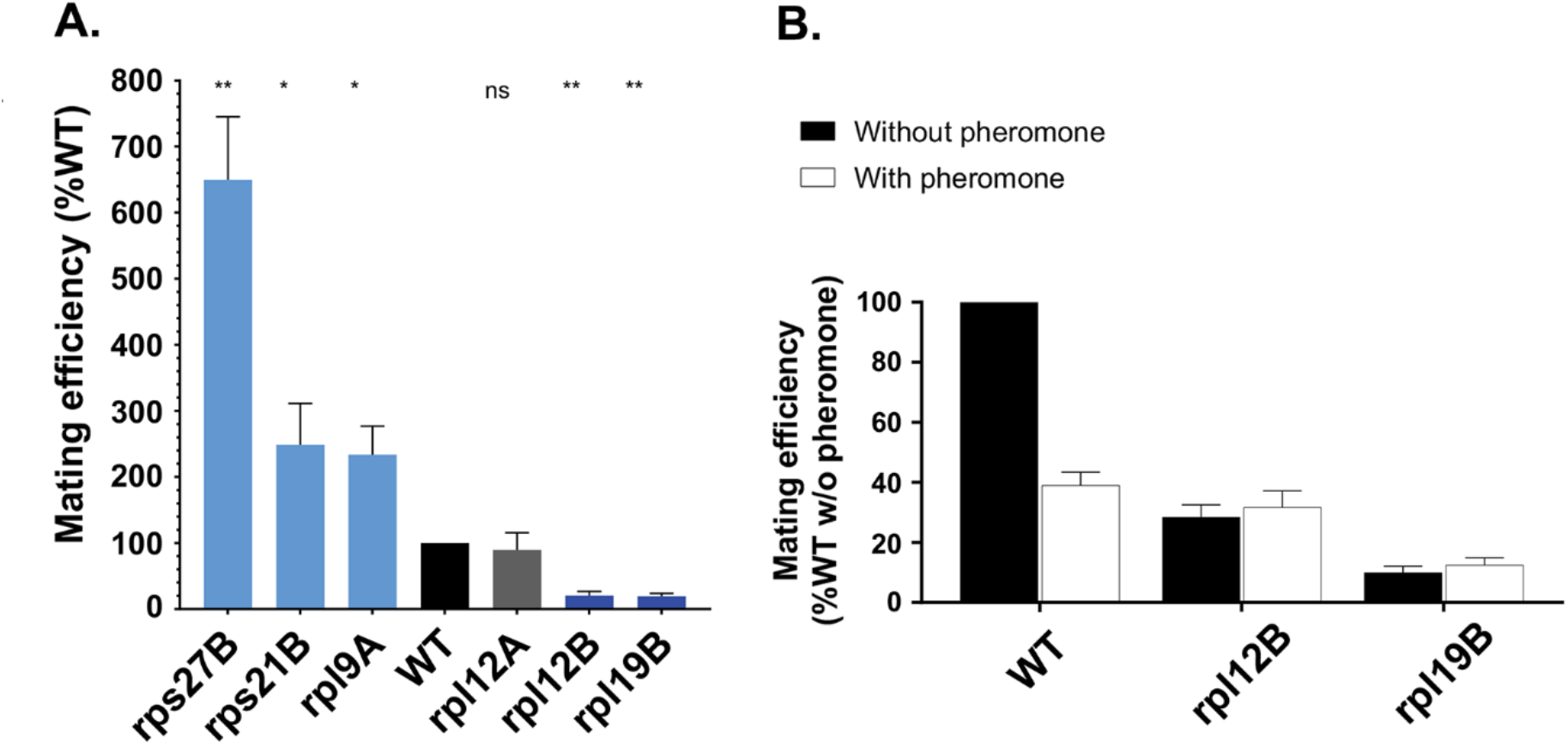
Validation of the ribosomal protein paralog mutants identified in the mating screen. (A) Quantitative mating assays were performed on a manually constructed set of ribosomal protein gene deletion mutants identified in the screen, and which showed positive and negative effects upon mating efficiency. The Y axis designates the % of mating obtained with control WT cells (BY4741) as 100%. The names listed on the X axis indicate the mutant genes tested. ns – not significant; * indicates *p* ≤0.05; **** indicates *p* ≤0.0001. (B) Confusion assays to assess mating efficiency were performed on ribosomal protein mutants as described in Figure 3.

### RP paralog specificity in pheromone signaling: *rpl12Δ* and *rpl19Δ* translatomes show reduced expression of proteins involved in mating

Recent work from our lab demonstrated that yeast lacking specific RP paralogs show large differences in their translatomes which incur changes in cell physiology^69^. This and other works suggest that RP paralog specificity in ribosomes is a defining element of translational control^70^. To examine the translatomes, we used PUNCH-P, which employs biotin-puromycin to label nascent polypeptides in isolated polysomes and mass spectrometry to identify translating proteins^69, 71^.

Using cells manually deleted for the paralogs, we examined the translatomes of *rpl12aΔ* vs. *rpl12bΔ* cells and *rpl19aΔ* vs. *rpl19bΔ* cells in comparison to WT cells grown on glucose-containing medium, and looked for the effects on proteins related to mating according to gene ontology (GO) terms (Figure 8A). We were able to identify >700 proteins for each strain, of which the translation of many mating/cytoskeletal-related proteins was altered in a paralog-specific manner (Figure 8B and C). Proteins with reduced translation in *rpl12bΔ* cells (relative to *rpl12aΔ* cells) and *rpl19bΔ* cells (relative to *rpl19aΔ* cells) included specific mating components (*e.g.* Ste6, Far1, and Hbt1), as well as Scp160, and proteins involved in actin regulation and endocytosis (*e.g.* Twf1, Abp1, Sac6, Myo2, Myo5, and Rsp5) (Figure 8B and C). In contrast, we observed the upregulation of translation of other endocytic and actin-related proteins in *rpl12bΔ* and *rpl19bΔ* cells, such as Rvs167, Srv2, and Bem2 (Figure 8B and C). Thus, the changes in translational control observed in *rpl12bΔ* and *rpl19bΔ* cells (Figure 8) appear to correlate with changes in chemotropism and mating (Figure 7).

**Figure 8.**
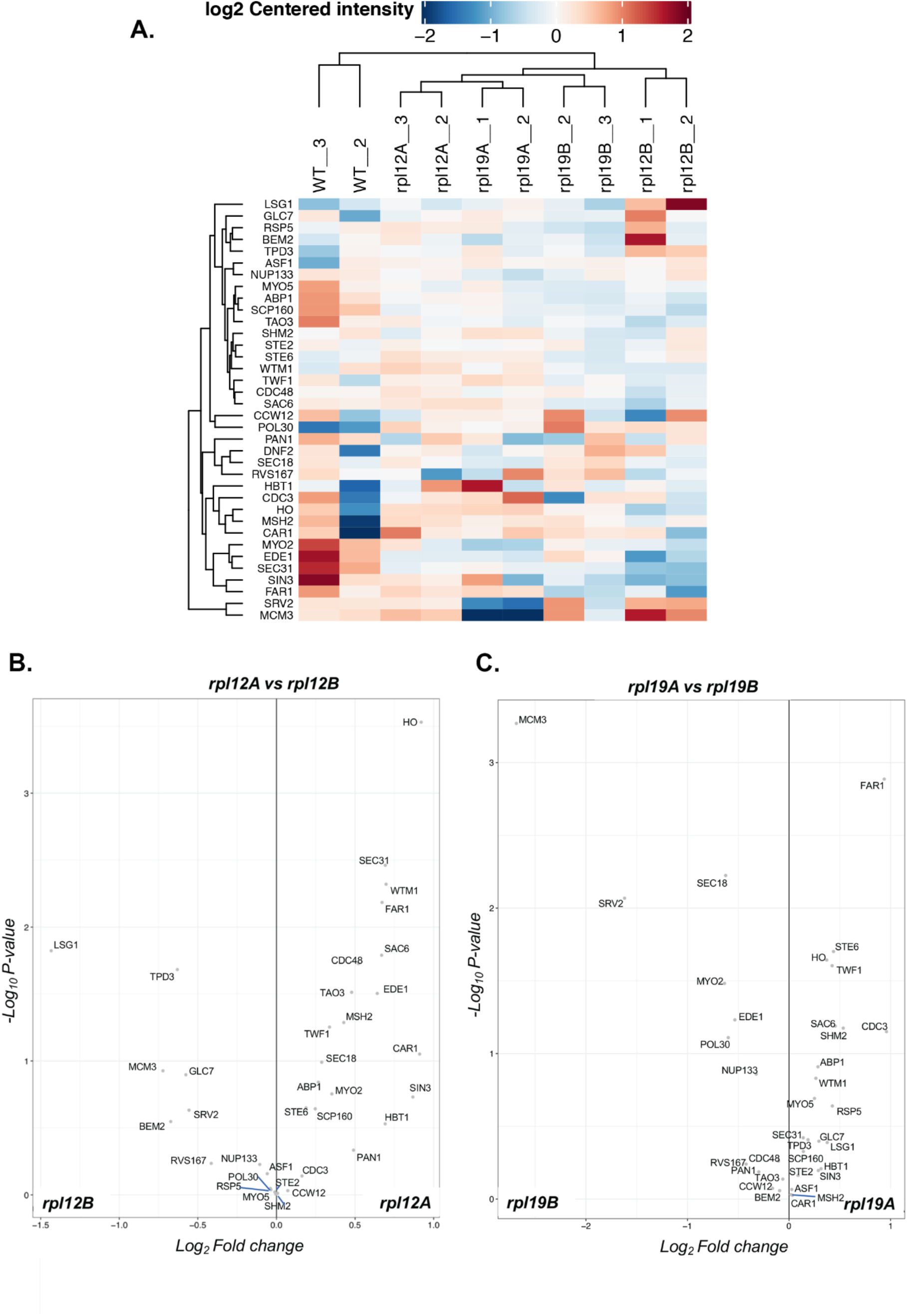
Ribosomal protein paralog specificity in the translation of mating pathway-associated proteins. (A) Translatome analysis of mating-related proteins in *RPL12* and *RPL19* paralog deletion mutants. WT, *rpl12aΔ, rpl12bΔ, rpl19aΔ,* and *rpl19bΔ* cells were grown to mid-log phase in rich medium (YPD) at 30°C, prior to polysome isolation and PUNCH-P analysis (see *Methods*). After mass spectrometry, hierarchal heatmap clusters of the mating-related translatome (*i.e.* nascent polypeptide chains) was performed. Note that the *rpl12bΔ* and *rpl19bΔ* mutants cluster separately from WT and their conjugate paralog mutants (*rpl12aΔ* and *rpl19aΔ*). (B and C) Volcano plots of the mating-related nascent proteins of *rpl12aΔ* vs *rpl12bΔ* (B) and *rpl19aΔ* vs *rpl19bΔ* (C).

## Discussion

The different MAPK cascades in yeast share conserved signaling modules containing kinases and scaffolding elements to generate intrinsic unidirectional signals that confer cellular responsiveness to a wide variety of extrinsic conditions (*e.g.* pheromone signaling, nutrient limitation, osmotic challenge, etc.)^72, 73^. Yet the properties that govern the interplay between the signaling pathways leading to MAPK activation and readout are not fully understood. While studies have identified mutations that block mating as a consequence of defects in pheromone signaling and morphogenesis^74, 75^, for example, no systematic attempt has been made to specifically identify regulators of chemotropism, a key feature of the cascade likely to be influenced by pathway cross- talk.

Here we have performed a systematic screen for positive and negative regulators of yeast mating, followed by secondary screening for mutants defective solely in gradient sensing and chemotropism (Figures 1-4 and 7, Table S1 and S2). Using this approach, we validated 13 genes that when mutated inhibit chemotropic growth and subsequent mating (Figures 2B and C, 3, 4 and 7), and 8 genes that act as negative regulators, as their mutation enhances mating (Figures 2C and D, 4, 7A, and Table S1). We initially classified these mutants into four different categories (Groups I-IV), depending upon their intrinsic levels of MAPK signaling, pheromone-induced morphogenesis, and angle of mating projection formation (a measure of gradient sensing) (Figure 4). Many of the regulators of chemotropism are known trafficking proteins involved in the delivery and retrieval of proteins to the cell surface, including the pheromone receptor, agglutinins, and pheromone. These include the exocyst genes *SEC3*, *SEC10*, *EXO84* and *SEC15*, along with other factors involved in protein transport and sorting, such as *YPP1*, *APS1*, *TRS3*1 and *BUL3*, which are all strongly defective in mating when mutated (Figure 3). Based upon their roles in exocytosis and endocytosis, these genes might be expected to affect mating *per se* (via defects in general protein transport). However, their impact upon the angle of incidence between mating partners (Figure 4C) suggests they also intimately affect gradient sensing. Since pheromone signaling rearranges the spatial distribution of the Ste2 pheromone receptor from the entire cell surface to the shmoo tip^76^, these mutants likely impinge upon receptor cycling and, hence, local signaling. This may also involve localization of a mobile polarity site (or patch) that serves as a site of communication where pheromones are released and sensed^77, 78^. Patch dynamics (via the delivery of secretory vesicles, endocytic recycling and actin dynamics) are thought to continually correct its position and align it with the gradient from the mating partner through local pheromone sensing^9^. The transmission of downstream signals through the polarity patch may involve genes connected to the regulation of MAPK signaling, such as *RAX1* (Group I), *BEM4* (Group II), and *AHK1* (Group III), identified in our screen (Figures 2A and B, and 3). These could potentiate patch dynamics, receptor signaling and cycling in part via phosphorylative control. Two of the Group IV genes, which are negative regulators of mating (Figure 2C and D), encode proteins that regulate the MAPK signaling module and buffer the pheromone signal. These include Gpr1 and Asc1, which are the receptor and a downstream effector for the nutrient sensing (filamentous growth) pathway, respectively^24, 25, 61, 62^, and compete with the pheromone receptors (Ste2 or Ste3) and Gpa1 for signaling via the Ste20-Ste11-Ste7 kinase cascade that activates the Kss1 MAPK, instead of Fus3^41^. Correspondingly, Hog1, a MAPK that operates in place of Fus3 downstream of the Ste20-Ste11-Pbs2 cascade and regulates high osmolarity sensing^64^, was also found as a negative regulator along with Msg5, which dephosphorylates Fus3^42, 43^. Thus, the inactivation of these genes potentiates the mating pathway presumably at the expense of signaling via the other MAPK paths. *IRA2* (also Group IV), which is a negative regulator of the RAS pathway that activates adenylyl cyclase and the nutrient sensing pathway^79^, was also found to be a negative regulator. In contrast, *AHK1*, which encodes a MAPK scaffolding protein that inhibits crosstalk to the filamentous growth pathway was found to be a potent positive regulator of mating. This likely results from its ability to restrict signaling along the Hog1 pathway that inhibits the mating signal^58^.

Finally, we identified RPs as specific regulators of the mating pathway, including positive (chemotropic growth) regulators, such as Rpl12b and Rpl19b, and negative regulators such as Rpl9a, Rps21b, and Rps27b (Figure 7) and Asc1 (Figures 2-6), which are involved in translational control. Thus, our screen essentially identified three sets of genes involved in: 1) protein trafficking; 2) MAPK signal transduction; and 3) translational control, as regulators of chemotropic growth and subsequent mating. A fourth set is composed of uncharacterized or poorly characterized proteins (*e.g*. negative regulators: *YGR016w*, *SYG1*, and *IRC19*; and positive regulators: *BUL3* and *SFD1*). How these proteins affect mating and chemotropism will require further research.

We note that we did not perform our confusion and chemotropism assays for all of our selected validated genes, nor many genes that were not chosen to be manually validated, including many RP paralog pairs. These will be studied in future research. Furthermore, it is likely that additional genes relevant to these processes were missed in our screen due to absence in the library, as a consequence of their impact upon mating during library preparation. Correspondingly, some DAmP library alleles may not yield a phenotype sufficiently robust to result in detectable defects in mating. And finally, our bias to remove genes associated with translation, mitochondrial function, and metabolism may have resulted in missed genes, as evidenced *post facto* (Figures 7 and 8).

As proteins involved in trafficking and signaling were obvious candidates for regulatory elements of mating and chemotropism, we were surprised to identify RPs as both negative (*e.g.* Asc1, Rpl9a, Rps21b, and Rps27b) and positive regulators (*e.g.* Rpl12b, and Rpl19b). Even though *ASC1* had been observed previously in a mating screen^33^, its participation in the mating process was never defined. While Scp160 has a known role in chemotropism^21^ and acts upon translational control along with Asc1^80^, it was unclear whether they co-regulate chemotropism. Here, we identified a tight genetic and physical interaction between Asc1 and Scp160 that likely explains their opposing effects on the mating and filamentous growth pathways. We found that the deletion of *ASC1*, which is required for filamentous growth^26^, led to large enhancement in mating efficiency, while its over-expression had the opposite effect (Figure S1). Moreover, Scp160 binds Asc1 upon pheromone stimulation or Gpa1 activation, but not upon Gpa2 activation (Figure 6). Deletion of *ASC1* led to increased mRNA binding by Scp160 (Figure 5), suggesting that Asc1 is a direct inhibitor of the RNA-binding function of Scp160. Even though chemotropism, mating, and downstream signaling (*i.e*. Fus1-GFP expression) are strongly and oppositely affected by deletion of *ASC1* and *SCP160* (Figures 2C and D, 3B, 4, and S1), and *FUS3* mRNA binding to Scp160 is altered in the absence of Asc1 (Figure 5), both proteins affect *FUS3* mRNA localization. Yet, this did not affect the level of Fus3 protein or phosphorylation (Figure S2). Thus, the effects exerted by the deletion of either gene on mating appear unconnected to altered *FUS3* mRNA localization. We suggest that the mating and filamentous growth MAPK pathways intersect at the level of Scp160 and Asc1, whereby pheromone/Gpa1-activated Scp160 mediates Asc1 binding to control readout of the two pathways at the level of translational control (Figure S3). Productive Scp160-Asc1 interactions, which have been shown to enhance translation on polysomes^29^, lead to propagation of the mating signal and, as a consequence, inhibition of the nutrient sensing/filamentous growth signals.

We also were surprised to see RPs as direct regulators of chemotropism and mating, which lends credence to the idea that incumbent with G*βγ* activation of Ste20 and downstream signaling, G*α* signals lead to changes in translational control of proteins connected directly to the mating pathway. This is borne out by experiments showing that the translatomes of *rpl12bΔ* and *rpl19bΔ* cells are deficient in proteins required for mating, as compared to WT, *rpl12aΔ* and *rpl19aΔ* cells (Figure 8). Thus, we predict that a translational control pathway involving Scp160 and specific RP paralogs plays a major role in propagation of the mating signal. While in contrast, Asc1 and other paralogs (*e.g.* Rpl9a, Rps21b, and Rps27b) may confer translational control of proteins involved in the propagation of filamentous growth pathway signals. Thus, it is tempting to implicate translational control as a direct means to modulate signals conferred by different MAPK cascades in order to control cellular responsiveness to extracellular signals. More work, including translatome analyses or ribosome profiling, is required to prove this point. In addition, these findings further extend our knowledge of RP paralog specificity, in terms of its direct involvement in translation and control of cell physiology^70, 81^.

Since gradient sensing and chemotropism/chemotaxis are highly relevant to the growth and migration of mammalian cells in complex microenvironments^82, 83^, there are likely to be structural and functional similarities to those underlying the yeast chemotropic response. Thus, some of the genes identified here may act as homologous regulators that participate in these processes. For example, mammalian Asc1 (RACK1) was earlier shown to regulate cell migration via its interaction with G*β*/*γ* and its downregulation enhances chemotaxis^84^. Numerous later studies reveal that this RP interacts with a wide variety of receptors and receptor-mediated signaling pathways to shuttle interacting partners, modify their activity, regulate their associations, and modulate stability^85^. Thus, Asc1/RACK1 is a prime candidate for MAPK pathway go-between to regulate chemotropic/chemotaxic responses to external signals. Whether its connection to mammalian Scp160 (vigilin)^86^ has been maintained in evolution remains unknown. Likewise, no study has examined the role of vigilin in either chemotropism or chemotaxis. Yet, our demonstration of Scp160 binding of Asc1 upon Gpa1 G*α* activation to downregulate its functions will prompt us in the future to determine the outcome upon cellular translation, and may prompt others to study their possible connection in higher organisms.

## Acknowledgements

The authors are grateful to Ralf Jansen (U. Tuebingen, Germany) for the gift of anti-Scp160 antibodies and to Pnina Weissman and Silvia Chuarzman for help with the screen in yeast. R.G-L. was supported in part by a returning scientist fellowship from the Israel Ministry of Absorption. This work was supported in part by an internal grants from the Jeanne and Joseph Nissim Center for Life Sciences Research and the Vice President’s Dedicated Equipment Fund, Weizmann Institute of Science. J.E.G. holds the Besen-Brender Chair in Microbiology and Parasitology, Weizmann Institute of Science. M.S is an incumbent of the Dr. Gilbert Omenn and Martha Darling Professorial Chair in Molecular Genetics. A.L., P.C. and R. G.-L. were supported in part by NIH grants GM084332 and CA209992.

## Methods

### Yeast Strains

Yeast strains used in this study are listed in Table S3.

### Mating efficiency screens

#### Automated library preparation

A SGA-ready *MATα* wild-type strain bearing the *TEF2pr-mCherry* gene integrated into the HO locus was crossed against 5119 yeast deletion strains^38^ (including 201 duplicate genes) and 1102 DAmP strains^22^ (Table S1 – Deletion & DAmP libraries) using the SGA method^87, 88^. A RoToR bench-top colony array instrument (pinning robot; Singer Instruments, Somerset UK) was used to pin the libraries. Strains from opposing mating types harboring the desired genomic manipulations (*i.e.* the mutation/deletion libraries on the one hand and the cytosolic mCherry WT strain on the other) were crossed and diploid cells selected on selective medium. Sporulation was induced by transferring yeast to nitrogen starvation media for 7 days, following which specific haploid progeny of *MAT*a were selected by growth on medium lacking histidine and using both canavanine and thia-lysine selection (Sigma-Aldrich) to select against remaining diploids. The resultant libraries contained the desired genotypes. Representative strains of the resulting yeast libraries were verified by PCR.

#### Mating efficiency and confusion screens

The query strain (*MATα* wild-type) expressing *TEF2pr-GFP*, as well as the *MAT*a gene knockout and attenuated (DAmP) libraries (384-well plates) were inoculated and grown in liquid YPD overnight at 30°C. Subsequently, cells were diluted and grown (4-5hrs) until reaching mid-log phase. Both the libraries and query strain were then plated by pinning (1.5mm diameter) onto separate YPD agar plates and grown overnight at 30°C. Following which, the *MATα* query strain and library plates were pinned together onto fresh YPD plates and allowed to grow for 4hrs at 30°C for mating. The individual *MATα*-*MAT*a crosses were then pinned into 1xTE in concanavalin-A coated 384-well glass bottom plates, centrifuged briefly to precipitate the cells, fixed using 4% paraformaldehyde (15min), and then washed twice with phosphate-sorbitol buffer (0.1M potassium phosphate pH7.5, 1.2M sorbitol). For the confusion assay, the *MATα*-*MAT*a crosses were plated in parallel onto fresh YPD agar plates or similar plates containing 30μM *α*-factor (Sigma-Aldrich) and allowed to grow for 4hrs at 30°C prior to fixation.

#### High-throughput microscopy

Crosses from the mating efficiency screen were visualized at room temperature using an automated inverted fluorescence microscopic ScanR system (Olympus Corp., Tokyo Japan). Images were acquired using a 60× air objective (NA 0.9) with excitation at 490/20 nm (GFP) or 572/35 nm (mCherry) and with an ORCA-ER charge-coupled device camera (Hamamatsu). After acquisition, images were manually reviewed and analyzed using the ImageJ analysis program.

#### IP and RNA binding

The immunoprecipitation of protein-RNA complexes was as described in Gelin-Licht *et al.* ^21^. Briefly, yeast cells expressing FLAG-tagged Scp160 were grown on synthetic medium to mid-log phase, lysed in lysis buffer (20mM Tris-Cl, pH 7.5, 150mM KCl, 5mM MgCl_2_, 100U/ml RNasin [Promega, Madison, WI, USA], 0.1% NP-40, protease inhibitor cocktail), and broken by vortexing with glass beads. For each anti-FLAG immunoprecipitation (IP) 4mg of the TCL was diluted in lysis buffer (final volume, 0.5ml), subjected to IP with anti-FLAG M2 affinity gel (30μl; Sigma, St. Louis, MO, USA) overnight at 4°C with rotation. The precipitates were then washed (x5) with lysis buffer. The precipitated protein-RNA complexes were eluted by incubation in 300μl of elution buffer, containing 50mM Tris-HCl, pH8.0, 100mM NaCl, 10mM EDTA, 0.1% SDS with 100μg/μl proteinase K (Promega, Madison, WI, USA), at 65°C for 30min.

RNA was purified from the IP eluates using the Masterpure^TM^ Yeast RNA purification kit (Epicenter, Madison, USA), according to the manufacturer’s instructions. After precipitation with added glycogen (8μg/ml; Roche, Mannheim, Germany) and 0.3M sodium acetate (pH5.0), the RNA was resuspended in 30μl of diethyl pyrocarbonate-treated water, treated with 3 units/sample of DNase I for 2 hours (Promega, Madison, WI, USA), and 1-3μl aliquots were taken for reverse transcription (RT) using the Moloney murine leukemia virus RT RNase H (−) (Promega, Madison, WI, USA), according to the manufacturer’s instructions, to yield cDNA. In parallel, total cellular RNA was purified from 15μl of TCL using the Masterpure^TM^ Yeast RNA purification kit, treated with DNase I, and 1-2μg of RNA was taken for RT, in the same manner as described above to yield total cDNA.

Real-time PCR was performed using the LightCycler 480 SYBR Green and LightCycler 480 system (Roche Diagnostics, Mannheim, Germany), according to the manufacturer’s instructions. Reaction mixtures (10μl final volume) contained the following components: 3μl of cDNA, 5pmol forward and reverse primers, and LightCycler 480 SYBR Green PCR Master mix, as prescribed by the manufacturer. The primer pairs for *FUS3* and *ASH1* produced only one amplicon when tested by conventional RT-PCR. The levels of specific mRNAs in the different strains tested were assessed relative to those obtained from precipitates derived from untreated (*i.e.* without pheromone or the expression of activated Gpa1) cells expressing FLAG-Scp160. The levels of specific mRNAs obtained from the different IP samples were normalized according to those of the total RNA.

#### Co-Immunoprecipitation

The immunoprecipitation of protein-protein complexes was as described in Gelin-Licht *et al.*^21^. Briefly, WT (BY4741) cells, WT cells expressing HA-tagged *ASC1* treated with or without *α*-factor, or HA-tagged *ASC1* cells co-expressing either *GPA1*^Q323**L**^ or *GPA2*^Q355L^ were grown to mid-log phase (O.D._600_ = ∼0.5) in 200ml synthetic selective medium at 26°C. Cultures were centrifuged at 3,000 x *g* for 5min at 4°C and resuspended in 1ml of lysis buffer (20mM Tris-Cl pH 7.5, 150mM KCl, 5mM MgCl2, 100U/ml RNasin, 0.1% NP-40, 1mM dithiothreitol, 2μg/ml aprotinin, 1μg/ml pepstatin, 0.5μg/ml leupeptin, and 0.01μg/ml benzamidine). Cells (100 O.D._600_ units each) were broken by vortexing with glass beads (0.5mm size) for 10min at 4°C and centrifuged at 10,000 × *g* for 10 min at 4°C to yield the total cell lysate (TCL). For IP, 0.5mg of TCL was diluted in lysis buffer (final volume, 1 ml) and subjected to IP with monoclonal anti-HA-Agarose antibody (A2095; Sigma) overnight at 4°C with rotation. Next, the precipitates were washed (×5) with lysis buffer and protein–protein complexes were eluted by incubation at 100°C for 5 min with an appropriate volume of 5X protein sample buffer (5X: 0.4 M Tris at pH 6.8, 50% glycerol, 10% SDS, 0.5 M DTT, and 0.25% bromophenol blue).

For Western analysis, samples were electrophoresed on 10% SDS-PAGE gels. Protein detection using anti-HA (H9658, Sigma) or anti-Scp160 antibodies (Gift of R. Jansen, U. Tuebingen) was performed using the Amersham ECL Western Blotting Detection Kit (GE Healthcare Life Sciences). Quantification of protein bands was performed using ImageJ software.

#### Quantitative Mating Assay

The quantitative mating assay was performed as described previously^89^. The assay is based on the complementation of auxotrophic markers present in *MAT*a and *MATα* cells upon crossing (*i.e. met15* mutation in *MAT*a BY4741 cells and *lys2* mutation in *MATα* BY4742 cells are rescued upon mating to yield a diploid capable of growing on medium lacking methionine or lysine). Cultures were grown to an OD_600_ = 0.3-0.4 at 26°C and 10^6^ cells of each strain (in a volume of ∼150μl), as well as *MAT*a and *MATα* mixtures, were collected by filtration on individual cellulose nitrate disk filters (25mm diameter, 0.45μm pore filters; BA-85 Schleicher & Schuell, Maidstone, UK) placed on top of the working filtration surface of a 500ml filtration apparatus (Corning, NY, USA). After filtration, each filter was placed cell-side up on YPD plates for 90min at 30°C, before resuspension in 1ml of SC-Met/-Lys (or SC-Trp/-Lys) in a sterile 15ml tube, followed by vortexing, serial dilution (1:10,100,1000), and plating 100μl of the dilutions onto SC-Met/-Lys (or SC-Trp/-Lys) plates to select for diploids. Mating efficiency was determined by colony counting after 3 days at 26°C.

#### Quantitative analysis of Fus1-GFP expression and shmoo formation

MAPK pathway activity and shmoo formation were measured by imaging flow cytometry (ImageStreamX Mark II, AMNIS-Luminex, Austin TX, USA). Yeast WT and mutant strains harboring a MAPK reporter (Fus1-GFP) were cultured in the presence of 5μM α-factor for 2hrs at 30°C, then fixed using 4% paraformaldehyde (15min), and washed twice with phosphate-sorbitol buffer. Cells were then analyzed by flow cytometry. Data was acquired using a 60X lens, and the lasers used were 488nm (100mW) and 785nm (5mW). Data was analyzed using the manufacturer’s software IDEAS 6.2 (AMNIS corp). Cells were gated according to GFP expression or according to size and ratio between length and width. .

#### Quantitative analysis of angle of zygote formation

Mutant and wild-type strains of mating type A, expressing mCherry from TEF2 promotor and wild type strain of mating type alpha, expressing GFP from TEF2 promotor were grown in YPD to mid-log phase. Equal numbers (1×10^6^) of *MAT*a and *MATα* were mixed and incubated for 4hr at 30°C. Cells were collected by filtration on individual cellulose nitrate filters (e.g. 25mm diameter, 0.45μm pore filters; BA-85 Schleicher & Schuell, Maidstone, UK) placed on top of the working filtration surface of a 500ml filtration apparatus (Corning, NY, USA). After filtration the cells were removed from the filters to TE medium and fixed with 4% paraformaldehyde.

Cells were imaged by an Imaging Flow Cytometer (ImageStreamX Mark II, AMNIS-Luminex, Austin TX, USA). Data was acquired using a 60X lens, and lasers used were 488nm (100mW), 561nm (200mW), and 785nm (5mW). Data was analyzed using the manufacturer’s software IDEAS 6.2 (AMNIS corp.). Images were compensated for spectral ovelap using single stained controls. Cells were first gated according to their area (in μm^2^) and aspect ratio (the Minor Axis divided by the Major Axis of the best-fit ellipse). Cells were further gated for focus using the Gradient RMS and contrast features (that measure the sharpness quality of an image by detecting large changes of pixel values in the image). Only objects with positive signals for both GFP and mCherry were gated, using single stained cells as controls. To identify the relevant cell interactions, two additional features were used: lobe count (that counts the lobes within an object) and Symmetry 3 (that measures the tendency of the object to have a three-fold axis of symmetry). To calculate the overlap between the GFP and mCherry, a similarity feature was used to measure the degree of which two images are linearly correlated, and was calculated as log transformed Pearson’s Correlation Coefficient.

### PUNCH-P

#### Nascent peptide Isolation

Nascent peptide Isolation and mass spectrometry for yeast were performed as previously described^69^. Briefly, 500 ml of yeast strains were grown in YPD at 30°C till mid-log phase (O.D._600_ = 0.6–0.8). Cells were washed once with ice-cold double-distilled water, snap-frozen, and stored at - 80°C. Frozen yeast cells were suspended in polysome extraction buffer (PEB; 20 mM Tris-HCl, pH 7.4, 140 mM KCl, 10 mM MgCl2, 0.5 mM DTT, 2 µg/ml leupeptin, 40 U/ml RNAsin, 1.4 µg/ml pepstatin, 0.2 mg/ml heparin [Sigma-Aldrich], EDTA-free complete protease inhibitor mix [Roche], and 1% Triton X-100) and lysed using glass beads (0.4–0.6 mm).The lysates were centrifuged at 17,400 x *g* for 30 min at 4°C to remove cell debris. Following this, 6ml of supernatant was loaded on a 2.5 ml 70% sucrose cushion. The samples were centrifuged at 60,000 rpm at 4°C for 2hr in an SW65 Ti ultracentrifuge rotor (Beckman Coulter, California, USA). The translucent polysome pellets were washed by gently dispensing and removing 500 µl of ice-cold RNase-free water. Ribosomes were suspended in 150 µl PEB (without detergent) snap-frozen and stored at -80°C. Purification of nascent polypeptide chains by Biotin-dc-puromycin labeling and pulldown using streptavidin beads was performed as described previous protocols^90^. For each sample, 12 O.D._254_ U of polysomes were incubated with 100 pmol of biotin-dc-puromycin (Jena Bioscience, Germany) per O.D._254_ unit at 37°C for 15 min. For each sample, 5 µl of streptavidin beads (GE Healthcare, Chicago, USA) per O.D._254_ unit was added, supplemented with 1 ml of SDS–urea buffer (50 mM Tris-HCl, pH 7.5, 8 M urea, 2% [wt/vol] SDS, and 200 mM NaCl), and incubated overnight at RT. The beads were washed five times with 1 ml SDS–urea buffer and incubated in 1 M NaCl for 30 min at 23°C followed by five washes in Ultrapure water.

#### Digestion and liquid chromatography mass spectrometry

The samples were subjected to on-bead tryptic digestion in the presence of urea. The resulting peptides were analyzed using nanoflow liquid chromatography (nanoAcquity; Waters Corp., Milford MA, USA) coupled to high resolution, high mass accuracy mass spectrometry. Each sample was analyzed on the instrument separately in a random order in discovery mode.

#### Data processing

Raw data was processed with MaxQuant v1.6.0.16. The data were searched using the Andromeda search engine against the *Saccharomyces cerevisiae* (strain S288c) proteome database appended with common lab protein contaminants and the following modifications: *e.g.* fixed modification - cysteine carbamidomethylation; variable modifications - methionine oxidation, asparagine and glutamine deamidation, and protein N-terminal acetylation. A total of 952 proteins were identified. Mating-related proteins were shorted by looking at the functional annotation of all identified proteins using DAVID^91^. We identified 34 out of 952 detected proteins as mating-related and used their translatome data for further analysis.

#### Differential protein expression analysis

Protein quantification was performed using the Differential Enrichment analysis of Proteomics data (DEP) package of R software (https://doi.org/doi:10.18129/B9.bioc.DEP). Two out of three biological replicates for each condition were selected for differential protein expression analysis. The intensity values were log-transformed and only proteins that had valid values in at least one experimental group were kept. Missing values were imputed (*i.e.* replacing missing values by a normal distribution of low values) ^92^. Finally, clustering (hierarchical) for each strain and volcano plots of one paralog vs. the other were performed using the same package.

---

**Figure S1.**
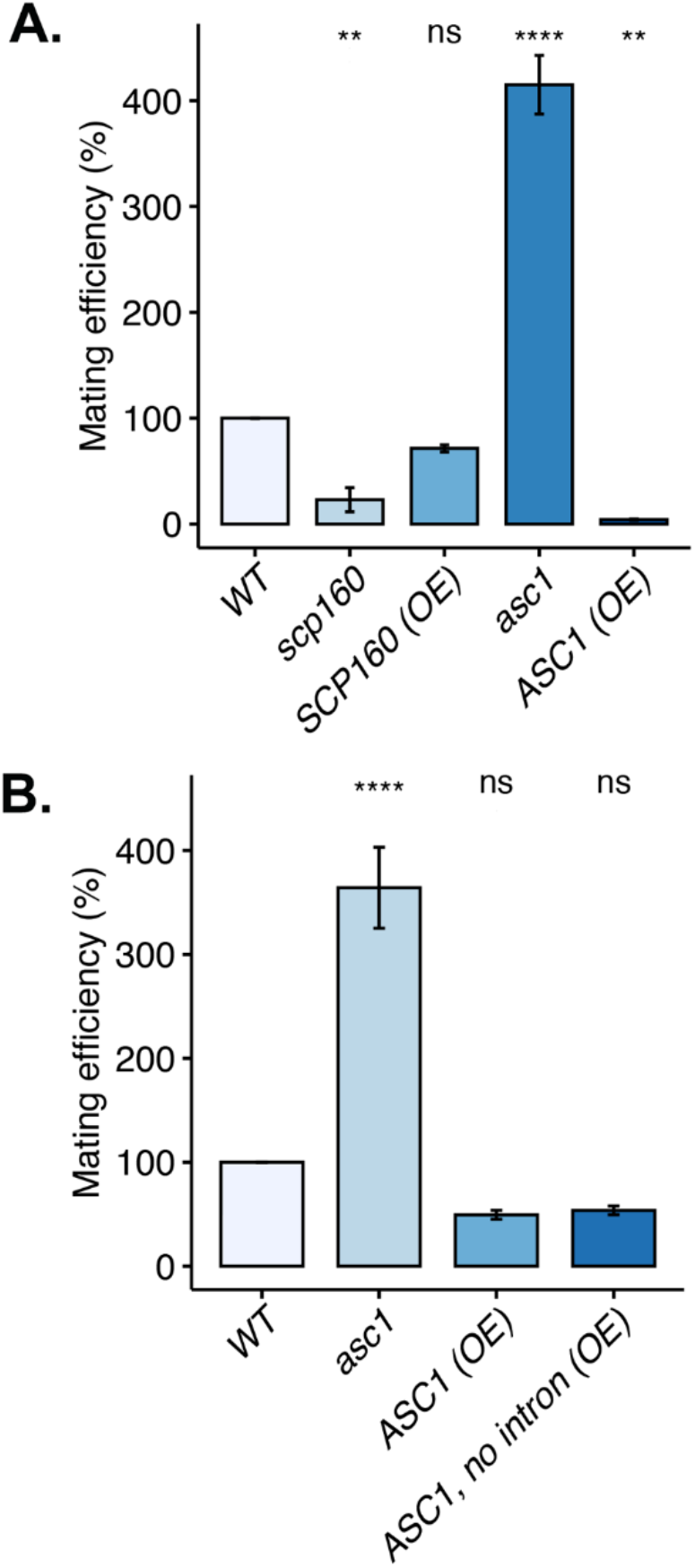
Asc1 and Scp160 modulate mating in an opposing fashion. (A) Over-expression or deletion of *ASC1* affects mating. Quantitative mating between wild-type (WT; BY4742 MATα cells) and *asc1Δ* (*MAT*a; *asc1*) cells, *scp160Δ* (*MAT*a; *scp160*) cells, and WT cells over-expressing *SCP160* (*SCP160*) or *ASC1* (*ASC1*) was compared to control BY4741 (*MAT*a) and BY4742 cells (WT). The mating efficiency of the WT control cells was designated as 100%. ns – not significant; ** indicates *p* ≤0.01; **** indicates *p* ≤0.0001. (B) Removal of the *ASC1* intron does not affect the ability of *ASC1* over-expression to inhibit mating. To test the necessity of the *ASC1* intron, we assayed quantitative mating between wild-type BY4742 (*MAT*α) cells and either *MAT*a cells over-expressing *ASC1* lacking its intron or native *ASC1*. Crosses between BY4742 (*MAT*α) cells and either *asc1Δ* or control WT cells were performed as controls. ns – not significant; **** indicates *p* ≤0.0001. °

**Figure S2.**
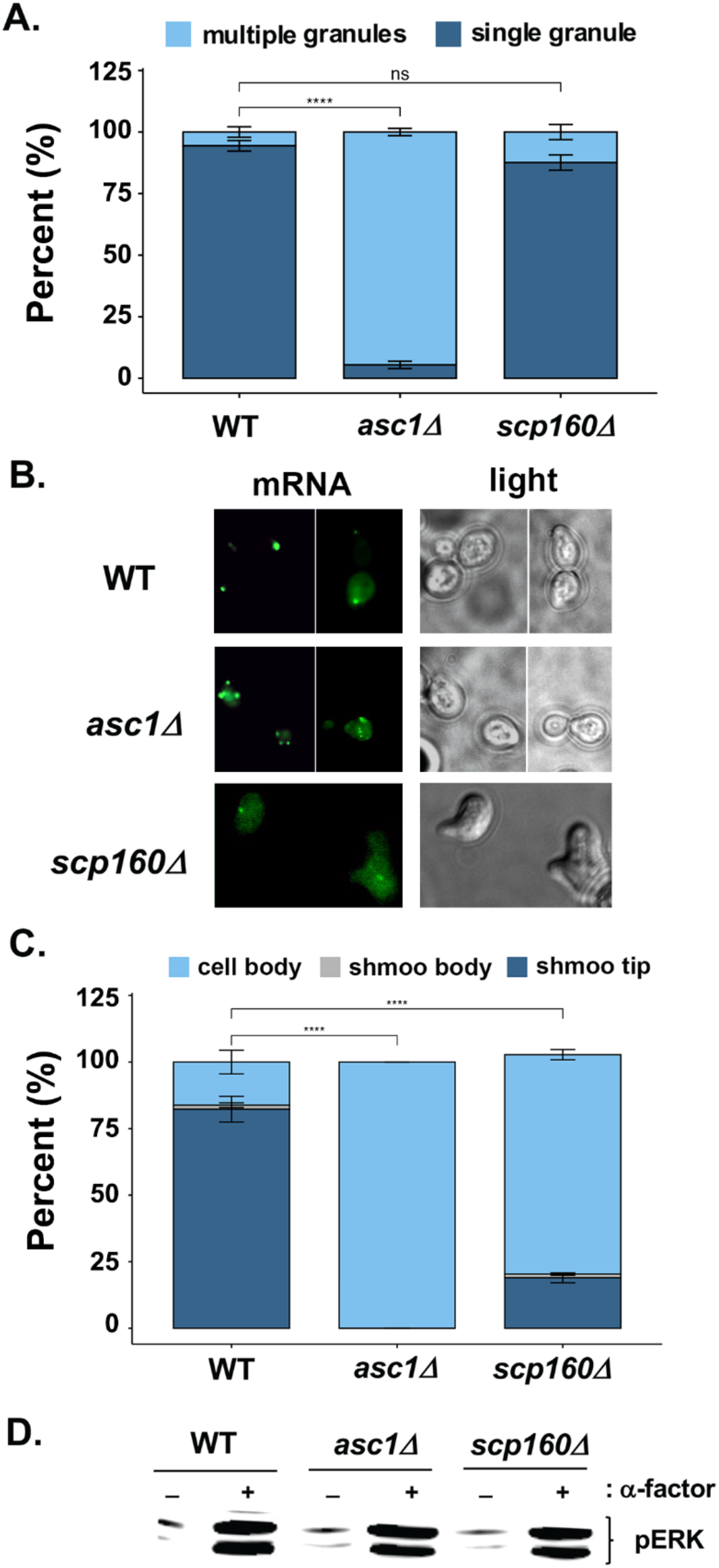
*FUS3* mRNA granule formation and localization in mating factor-treated yeast cells is controlled by Scp160 and Asc1. (A) Multiple *FUS3* granules accumulate in *asc1Δ* cells treated with α-factor. *MAT*a wild-type (WT), *asc1Δ* cells, and *scp1601Δ* cells endogenously expressing MS2 aptamer-tagged *FUS3* mRNA and co-expressing MS2-CP-GFP(x3) were grown at 30°C on synthetic medium and treated for 2hrs in the presence of 5μM α-factor, prior to scoring for shmooing cells with visible RNA granules. Cells with either single (dark blue bar) or multiple (light blue bar) granules were scored using fluorescence microscopy. ns – not significant; **** indicates *p* ≤0.0001. (B) *FUS3* mRNA granules are mislocalized in both *asc1Δ* cells and *scp1601Δ* cells. The mRNA granules scored in (*A*) were further examined for their intracellular localization, either to the cell body (soma; light blue) shmoo body (base; grey), or shmoo tip (dark blue). Representative images are shown. (C) *FUS3* mRNA granules are mislocalized to the cell body in both *asc1Δ* cells and *scp1601Δ* cells. Quantification of *FUS3* mRNA localization in the cells scored in (*A*). ns – not significant; **** indicates *p* ≤0.0001. (D) Phosphorylated Fus3 protein levels are unchanged in the absence of Asc1 or Scp160. BY4741 wild-type (WT), *asc1Δ*, and *scp160Δ* cells were grown to mid-log phase and treated with 5μM α-factor for 1hr to induce *FUS3* expression. Cells were harvested and lysed for whole cell protein extraction. The lysate was separated on a 10% gradient SDS-PAGE gel and analyzed by immunoblotting with anti-pERK antibody, which detects both phosphorylated Fus3 (upper band) and Kss1 (lower band).

**Figure S3.**
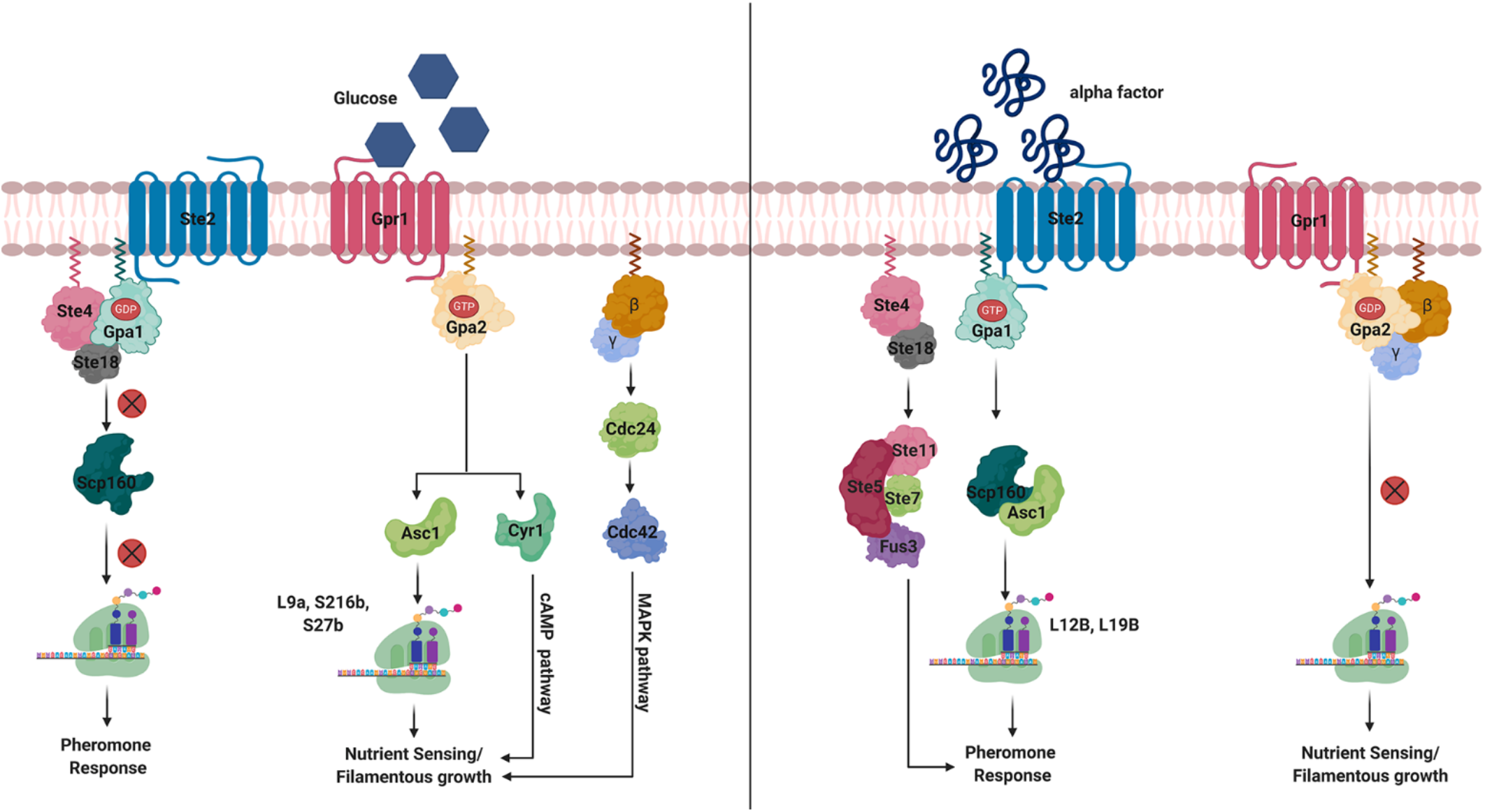
A model for translational control of the pheromone- and nutrient-sensing/filamentous growth signaling pathways during the mating response in yeast. Left panel: Glucose stimulation of the Gpr1 GPCR leads to signaling via the Gpa2 G*α* subunit to activate Asc1 and Cyr1 (adenylyl cyclase)

**Table S3.** Yeast strains

| Strain | Genotype | Source |
| --- | --- | --- |
| BY4741 | *MATa his3Δ1 leu2Δ0 met15Δ0 ura3Δ0* | Euroscarf |
| BY4742 | *MATα his3Δ1 leu2Δ0 met15Δ0 ura3Δ0* | Euroscarf |
| *scp160Δ* | *MATa his3Δ1 leu2Δ0 met15Δ0 ura3Δ0 scp160Δ::kanMX* | Euroscarf |
| JFy4493 (W303) | *MATa ade2-1 his3 leu2-3 trp1-1 ura3-1 FLAG-SCP160::kanMX4* | J. Fridovich-Keil |
| JFy5335 | *MATa ade2-1 his3 leu2-3 trp1-1 ura3-1 FLAG-SCP160KHΔ14::kanMX4* | J. Fridovich-Keil |
| JFy4493 *asc1Δ* (W303) | *MATa ade2-1 his3 leu2-3 trp1-1 FLAG-SCP160::kanMX4 asc1Δ::URA3* | This study |
| *ASC1-HA* (BY4741) | *MATa his3Δ1 leu2Δ0 met15Δ0 ura3Δ0 ASC1-HA::NAT* | This study |
| *ASC1-HA* (JFy4493) | *MATa ade2-1 his3 leu2-3 trp1-1 ura3-1 FLAG-SCP160::kanMX4 ASC1-HA::NAT* | This study |
| TEF2-Cherry SGA (S288C) | *MATα ura3Δ0::TEF2pr-Cherry::URA3 his3Δ1 leu2Δ0 met15Δ0 can1Δ::STE2pr-spHIS5 lyp1Δ::STE3pr-LEU2* | M. Schuldiner |
| TEF2-GFP SGA (S288C) | *MATα his3Δ1 leu2Δ0 met15Δ0 ura3Δ0 can1Δ::STE2pr-sp HIS5 lyp1Δ::STE3pr-LEU2 TEF2pr-GFP::NAT* | M. Schuldiner |
| Gene X-NAT<sup>1</sup> (BY4741) | *MATa his3Δ1 leu2Δ0 met15Δ0 ura3Δ0 X::NAT* | This study |
| FUS1-GFP (BY4741) | *MATa his3Δ1 leu2Δ0 met15Δ0 FUS1-GFP::URA3* | This study |
| Gene X-NAT FUS1-GFP<sup>2</sup> (BY4741) | *MATa his3Δ1 leu2Δ0 met15Δ0 FUS1-GFP::URA3 X::NAT* | This study |

<sup>1</sup>The genotype of the 52 knockout strains used for manually validation (Table S1; sheet - *Manual validation*)
<sup>2</sup>The genotype of the wild-type and knockout mutant strains expressing *FUS1* reporter for the analysis of the MAPK pathway activity (Figure 4A)

